# Generation of Human Taste Bud Organoids as a Human-Mimetic Platform for Modeling Taste Perception

**DOI:** 10.64898/2026.08.09.743674

**Authors:** Jiyeon Chae, Soon Sung Kwon, Jisoo Kim, Hyunjin Moon, Van Quan Do, Sofie Zehentner, Hyung-Ju Cho, Jinhyuk Bhin, Seok Jun Moon, Chul Hoon Kim

**Author notes:** These authors contributed equally to this work (Jiyeon Chae, Soon Sung Kwon, Jisoo Kim).

## Abstract

We established a human taste bud organoid system derived from circumvallate papillae. This model has been highly anticipated in the field of taste research, where feasible approaches for validating taste biology discovered in rodent models have been limited. Through a stepwise exploratory strategy, we systematically identified and optimized the niche factors required to maintain taste bud organoids and promote their differentiation. This human taste bud organoid system comprises Type I–IV taste receptor cells (TRCs) as well as stem/progenitor cells, and its sensory receptor cells exhibit calcium responses to taste stimuli. Using this system, we identified robust Wnt signaling as a requirement for optimal TRC fate progression, uncovered a human-specific transcriptional program in LGR5⁺ cells, and identified previously unrecognized molecular markers for Type I TRCs. By recapitulating native human taste bud cell diversity and function, this organoid provides a tractable platform for studying human taste biology and dysfunction.

## Introduction

Taste is a fundamental sensory modality that enables organisms to evaluate the nutritional value and potential toxicity of ingested substances^1,2^. As such, the gustatory system plays a central role in regulating feeding behavior and maintaining metabolic health across species^2–4^. In humans, taste extends beyond its physiological functions, contributing to social interactions and quality of life. Accordingly, taste dysfunction associated with cancer therapies^5^, viral infections including COVID-19^6^, aging^7^, and other clinical conditions^8^ can substantially impair nutrition and overall health outcomes, with broader consequences for social participation and overall well-being.

Taste buds are specialized multicellular sensory organs embedded in the oral epithelium that detect the five canonical taste qualities — sweet, umami, bitter, sour, and salty^9,10^. Mature taste buds comprise functionally distinct principal cell types^10^. Glia-like Type I cells provide structural and homeostatic support^11,12^ and participate in sweet adaptation^13^. Type II cells mediate sweet, umami, and bitter taste detection through T1R and T2R G protein–coupled receptors and canonical PLCβ2 signaling^11,14^ and also mediate appetitive sodium taste by ENaC in a distinct subset of Type II cells that lack TRPM5 and gustducin^15^. Type III cells transduce sour stimuli through the OTOP1 proton channel^16,17^. These three taste receptor cell (TRC) subtypes are short-lived and undergo continuous renewal throughout life, with an average turnover period of approximately 10–14 days^18^, making taste buds one of the most rapidly renewing peripheral sensory tissues in mammals. In mouse taste buds, this renewal is sustained by basal stem/progenitor cells, which ultimately give rise to post-mitotic Type IV precursors expressing sonic hedgehog (SHH)^19,20^. These precursors subsequently differentiate into mature Type I–III taste cell subtypes^19,20^.

Much of our current understanding of taste stem cell biology derives from mouse models, where Lgr5 marks an actively cycling, Wnt-responsive stem/progenitor population within the Krt5/Krt14-expressing basal epithelium of the circumvallate and foliate papillae^21,22^. These cells give rise to all types of mature TRCs under the control of microenvironment-derived signaling cues^23–25^. However, whether similar self-renewal and differentiation programs operate in human taste buds remains largely undefined. Direct studies of native human taste tissue are constrained by the scarcity of available samples, creating a need for a scalable and sustainable human taste bud cell culture system. Such a system would enable investigation of the cellular and molecular mechanisms underlying human taste bud maintenance and renewal, as well as the organization of the human taste receptor repertoire and associated signal transduction pathways.

Adult stem cell-derived organoids have created new opportunities for studying epithelial biology by enabling the long-term maintenance and expansion of epithelial stem/progenitor cells, promoting their differentiation into epithelial lineages, and facilitating functional modeling across diverse epithelial tissues, including the intestine, stomach, and liver^26–29^. Taste bud organoids have likewise been generated from adult stem/progenitor cells in mice, pigs, and non-human primates^30–32^. A human taste organoid, however, has remained elusive, implying that the maintenance and differentiation of human taste lineages require species-specific culture conditions that cannot be readily recapitulated through the simple adaptation of protocols established in other organisms.

Here, we report a human taste bud organoid (hTBO) system derived from adult human circumvallate papillae (hCVPs), thereby providing a human cell-based platform that substantially advances the experimental study of human taste biology. Through systematic optimization of niche signals, we established a 2-phase expansion-to-differentiation protocol that generates hTBOs faithfully recapitulating the cellular composition and molecular identity of native human taste buds, and revealed a previously unrecognized dependence of human TRC fate progression on heightened Wnt/R-spondin signaling. Moreover, single-cell transcriptomic profiling resolved a stepwise gustatory differentiation trajectory, progressing from basal and cycling basal cells through highly Wnt-responsive transitional basal and LGR5⁺ cells to post-mitotic precursors and mature TRCs. Cross-species analysis revealed divergent gene regulatory programs in LGR5⁺ cells and identified novel Type I TRC markers. Functionally, hTBOs contained distinct tastant-responsive cell populations and accurately reproduced *TAS2R38* haplotype-dependent responses to phenylthiocarbamide (PTC). Collectively, this work uncovers human-specific features of taste bud cells and establishes a tractable platform for future studies of human taste biology and taste disorders.

## Results

### Establishment of a long-term expandable hTBO

To establish an hTBO platform, we isolated hCVP tissue and dissociated it into single cells, which were then embedded in Matrigel domes for systematic screening of culture conditions that support long-term organoid expansion and maintenance *in vitro* (Figure 1A). Because the signaling requirements for maintaining hTBOs had not yet been defined, we initially adopted a “complete” organoid medium containing a broad range of factors previously validated in other epithelial organoid systems for their ability to support epithelial stem cell maintenance and long-term expansion^27–30^. Under these conditions, dissociated hCVP cells reproducibly formed three-dimensional epithelial organoids (Figures 1A-1D). To define the minimal signaling requirements for long-term proliferation and maintenance, we performed systematic single factor-withdrawal experiments. Removal of Wnt3a-conditioned media (Wnt3a CM), R-spondin 1, Noggin, A83-01 (a TGFβ pathway inhibitor), FGF10, or forskolin (a cAMP activator) markedly reduced culture longevity and expansion capacity, indicating that Wnt activation, BMP inhibition, TGFβ suppression, FGF signaling, and cAMP elevation are each required to sustain hTBO growth (Figure 1B). In contrast, removal of EGF, nicotinamide, or SB202190 (a p38 MAPK inhibitor) did not significantly affect organoid viability, long-term maintenance (Figure 1B), or continued expansion through multiple serial passages across three independent donor-derived lines (Figure 1C). Together, these data define a minimal set of signaling factors—Wnt3a, R-spondin 1, Noggin, TGFβ inhibitor, FGF10, and forskolin (hereafter, the “essential medium”)—that supports robust long-term maintenance of hTBOs.

**Figure 1.**
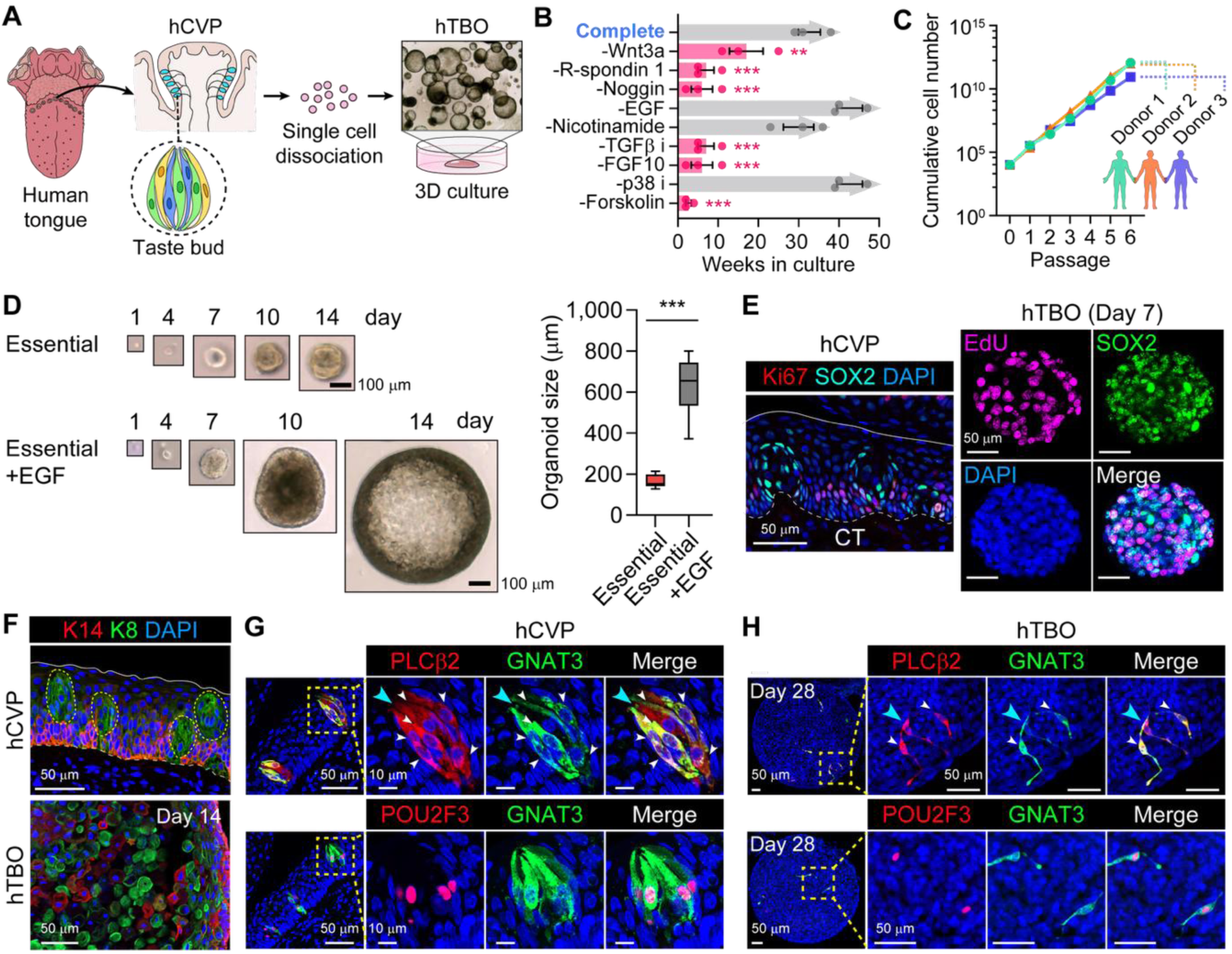
Establishment of long-term expandable human taste bud organoid (hTBO). (A) Schematic of hTBO generation from human circumvallate papilla (hCVP) tissue. Single epithelial cells isolated from hCVP were embedded in Matrigel, where they gave rise to 3D organoids. (B) Single-factor withdrawal analysis defining the signaling requirements for long-term hTBO maintenance (n = 3 biological replicates from three independent donors). Arrowheads denote conditions under which organoids continued to expand actively at the final time point analyzed. \*\**P* < 0.01 and \*\*\**P* < 0.001 compared with Complete; one-way ANOVA followed by post hoc Bonferroni correction. TGFβ i, TGFβ inhibitor (A83-01); p38 i, p38 MAPK inhibitor (SB202190). (C) Cumulative cell expansion of hTBOs maintained in essential medium over serial passaging in three independent donor-derived lines (Donors 1–3). (D) Representative bright-field images showing the growth of single-cell-derived organoids cultured with or without EGF over time (left), and quantification of organoid diameter under each condition at day 14 (right; n = 9 independent organoids from a single donor). \*\*\**P* < 0.001 compared with Essential medium; two-tailed unpaired Student’s *t* test. Scale bars, 100 μm. (E) Immunostaining of hCVP tissue and hTBOs for the proliferation markers Ki67 and EdU and the progenitor-associated marker SOX2. Solid and dashed lines indicate the apical and basal surfaces, respectively. CT, connective tissue. Scale bars, 50 μm. (F) Immunostaining for the basal cell marker KRT14 and the taste cell marker KRT8 in hCVP tissue and hTBOs at culture day 14. White solid and dashed lines indicate the apical and basal surfaces, respectively. Taste buds are outlined by yellow dashed ovals. Scale bars, 50 μm. (G) Validation of antibodies against Type II TRC markers in hCVP tissue. Representative images show co-staining of PLCβ2 and GNAT3 (top) or POU2F3 and GNAT3 (bottom). Boxed regions are shown at higher magnification. White arrowheads mark PLCβ2/GNAT3 co-expressing cells. A cyan arrowhead indicates a PLCβ2 single-positive cell. Scale bars, 10 μm and 50 μm. (H) Detection of differentiated Type II TRC markers in hTBOs at culture day 28. Representative images show expression of PLCβ2 and GNAT3 (top) or POU2F3 and GNAT3 (bottom). Boxed regions are shown at higher magnification. White arrowheads mark PLCβ2/GNAT3 co-expressing cells. A cyan arrowhead indicates a PLCβ2 single-positive cell. Scale bars, 50 μm. Data are presented as means ± SEM.

Despite stable long-term maintenance in the essential medium, hTBOs remained relatively small, attaining only approximately 100 µm in diameter by day 14 and consequently providing limited cell numbers for downstream applications (Figure 1D). We therefore asked whether reintroducing EGF into the essential medium could enhance organoid growth. Indeed, EGF supplementation increased hTBO diameter to approximately 700 µm by day 14 before growth plateaued (Figures 1D and S1A). Thus, we elected to retain EGF in the culture medium to enhance organoid biomass for downstream applications.

We next asked whether hTBOs recapitulate the major cellular populations of the native hCVP epithelium, including intragemmal cells and extragemmal populations such as non-gustatory epithelial cells and basal progenitors. Proliferating cells were detected in hCVP tissue by Ki67 immunostaining and in hTBOs by EdU incorporation (Figure 1E). In hCVP tissue, Ki67^+^ cells were localized to the basal epithelial region, consistent with the known distribution of basal progenitors in mouse taste bud tissue^23^. In contrast, EdU⁺ cells in hTBOs lacked a discernible spatial organization, suggesting that hTBOs do not fully recapitulate the architectural features of native tissue. Nevertheless, hTBOs clearly harbored a robust proliferative cell population sufficient to support continued organoid growth *in vitro*. SOX2^+^ stem/progenitor cells have been shown to contribute actively to taste bud cell production in mice^33,34^. In hCVP tissue, SOX2 was broadly expressed in the lingual epithelium but exhibited stronger expression in and around taste buds than in the surrounding basal epithelium, with a subset of SOX2⁺ cells co-expressing Ki67 (Figure 1E, left). This pattern is consistent with previous findings that high SOX2 expression predicts taste lineage competency in lingual progenitors^35^. In hTBOs, a subset of EdU^+^ cells also expressed SOX2 (Figure 1E, right). Together, these findings suggest that an actively proliferating SOX2^+^ stem/progenitor-like population is present in hCVP tissue and is recapitulated in the corresponding hTBOs. Using KRT14- and KRT8-specific antibodies, we further found that KRT14^+^ cells were localized to the basal region, whereas KRT8^+^ cells were confined to intragemmal regions in human taste bud tissue, consistent with marker distributions previously reported in mouse taste buds^22,23^. Guided by this validation of antibodies in native tissue, we found that hTBOs similarly contained KRT8^+^ post-mitotic intragemmal cells and KRT14^+^ basal progenitor cells with minimal overlap (Figure 1F), mirroring the basal-to-taste cell organization observed in native tissue. A KRT8^−^/KRT14^−^ or KRT8-low/KRT14-low cell population was also detected in hTBOs (Figure 1F), consistent with the presence of non-gustatory epithelial cells surrounding taste buds in native tongue tissue^23^.

To determine whether hTBOs also generate fully specified TRCs, including Type I–III cells, we examined canonical markers of mature taste cell subtypes that have been validated in mouse taste buds. Screening of reported antibodies revealed that many failed to produce specific staining in human taste bud tissue. Only a subset of the tested antibodies against the Type II TRC markers POU class 2 homeobox 3 (POU2F3), phospholipase Cβ2 (PLCβ2), and G protein subunit alpha transducin 3 (GNAT3), showed specific staining patterns in hCVP tissue (Figure 1G). Using these validated antibodies, we detected sparse POU2F3⁺, PLCβ2⁺, and GNAT3⁺ cells in hTBOs, providing an early indication that the human organoid system supports differentiation into TRCs, at least of the Type II lineage (Figure 1H).

### Optimization of hTBO culture for TRC differentiation

Growth factor-rich culture conditions have been widely used in various human adult stem cell-derived organoid systems to support long-term expansion and maintenance^27–29,36,37^ and in many cases, differentiation as well. Mouse taste bud organoids (mTBOs) proliferate robustly and readily differentiate into TRCs in medium containing key mitogenic cues and Wnt-supporting factors, including EGF and R-spondin 1^30,38,39^. However, when applied to hTBOs, a comparable condition consisting of essential medium supplemented with EGF (hereafter referred to as the “1-phase culture method”) yielded only sparse mature TRCs by 28 days *in vitro* (Figure 1H), substantially limiting the utility of this human organoid model. Because terminal differentiation is generally coupled to cell-cycle exit, whereas cell-fate transitions frequently occur in conjunction with cell-cycle progression in proliferating cells^40^, we reasoned that sustained mitogen-driven proliferative signaling might impede TRC maturation. Indeed, mature TRCs are generally post-mitotic, whereas basal stem/progenitor cells remain proliferative and reside within a distinct anatomical and signaling niche that promotes self-renewal and preserves their competence for subsequent differentiation into mature taste cells in mouse taste bud tissue^22^. However, unlike in native taste bud tissue, organoid cultures subject these different cell populations distinguished by their cell-cycle states, proliferative stem/progenitor cells and differentiating cells, to the same signaling environment. To address this inherent limitation of organoid culture, we established a phase-dependent stimulation strategy. We first withdrew EGF, a mitogenic factor previously included to support expansion, after a 7-day expansion phase to determine whether its removal would promote lineage commitment and terminal TRC maturation. Consistent with this idea, withdrawal of EGF from the 1-phase culture condition increased Type II TRC numbers, although organoid growth was concomitantly reduced (Figure S1B). To further reduce exogenous mitogenic input, we additionally considered withdrawal of Wnt3a CM (50%). Wnt3a-CM is conventionally produced in FBS-containing basal medium^41^, and FBS contains a variety of biologically active components, including growth-promoting and mitogenic factors^42^. Guided by these considerations, we built the framework for a 2-phase culture strategy where hTBOs were cultured in expansion media for 7 days to accumulate biomass and then shifted to differentiated medium lacking EGF and Wnt3a CM, thereby reducing mitogenic input and promoting TRC differentiation (Figure 2A). Instead of Wnt3a CM as a source of Wnt ligand, we used the GSK3 inhibitor CHIR99021 to activate the canonical Wnt activation pathway in this framework (Figure 2A). Titration of CHIR99021 showed that Type II TRC marker-positive cells increased up to 3 μM, whereas higher concentrations failed to further enhance differentiation and reduced organoid size (Figure S2A). We therefore used 3 μM CHIR99021 for subsequent experiments and developed the foundational protocol for the 2-phase culture method (Figure 2A). Compared with both the 1-phase culture method and an intermediate 2-phase condition lacking only EGF, the optimized 2-phase culture method markedly increased the abundance of POU2F3-, PLCβ2-, and GNAT3-expressing Type II TRCs (Figures 2B and 2D), demonstrating that temporal separation of a Wnt-stimulated differentiation phase from a mitogen-rich expansion phase serves as an effective strategy for enhancing TRC differentiation in hCVP-derived organoids.

**Figure 2.**
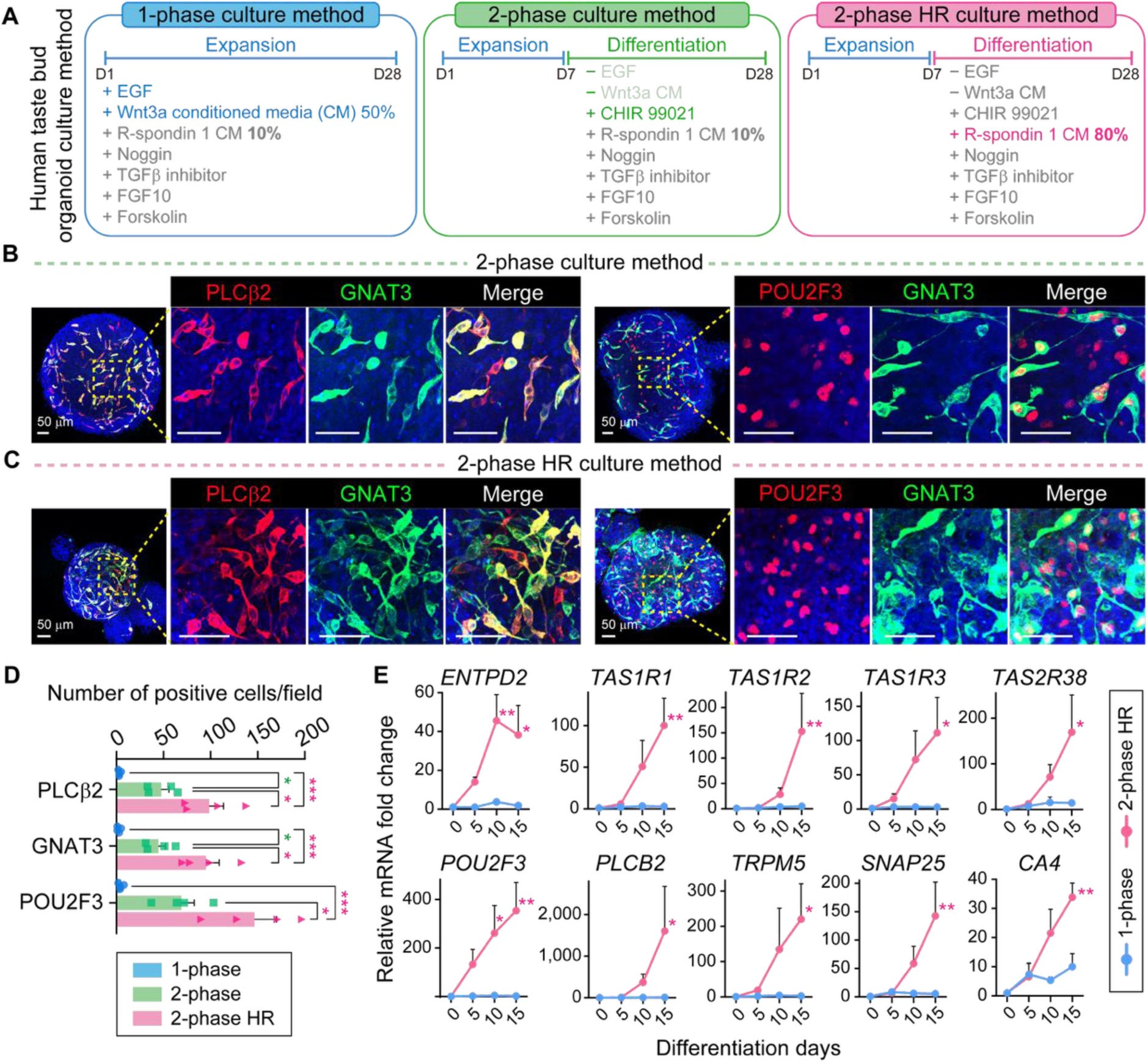
Optimization of hTBO culture conditions for enhanced TRC differentiation. (A) Schematic of the three culture conditions used to optimize TRC differentiation from hTBOs. In the 1-phase culture method, organoids were maintained in expansion medium throughout. In the 2-phase culture method, organoids were expanded for 7 days and then switched to differentiation medium lacking EGF and Wnt3a-CM, with CHIR99021 substituted as a defined Wnt agonist. In the 2-phase high R-spondin (HR) culture method, organoids were transitioned to the same differentiation conditions, except that R-spondin 1 CM was increased from 10% to 80% during the differentiation phase. (B) Representative immunofluorescence images of hTBOs cultured using the 2-phase method and stained for the Type II TRC markers PLCβ2 and GNAT3 (left) or POU2F3 and GNAT3 (right). Boxed regions are shown at higher magnification. Scale bars, 50 μm. (C) Representative immunofluorescence images of hTBOs cultured using the 2-phase HR method and stained for PLCβ2 and GNAT3 (left) or POU2F3 and GNAT3 (right). Boxed regions are shown at higher magnification. Scale bars, 50 μm. (D) Quantification of PLCβ2^+^, GNAT3^+^, and POU2F3^+^ cells per field of view under the indicated culture conditions (n = 4 independent organoid cultures from three independent donors). Each field of view represents an area of 300 μm × 300 μm. \**P* < 0.05 and \*\*\**P* < 0.001 compared with 1-phase; one-way ANOVA followed by post hoc Bonferroni correction. (E) Longitudinal qRT-PCR analysis of TRC differentiation markers during the differentiation phase under the indicated culture conditions, showing relative mRNA fold changes over time for Type I (*ENTPD2*), Type II (*TAS1R1*, *TAS1R2*, *TAS1R3*, *TAS2R38*, *POU2F3*, *PLCB2*, and *TRPM5*), and Type III (*SNAP25* and *CA4*) TRC marker genes (n = 3 independent organoid cultures from three independent donors). \**P* < 0.05 and \*\**P* < 0.01 compared with 1-phase; two-way ANOVA followed by post hoc Šídák correction. Data are presented as mean ± SEM.

Our 2-phase culture experiments demonstrated that Wnt signaling drives TRC differentiation in hTBOs. In particular, the finding that increasing CHIR99021 from 0 to 3 μM promoted TRC differentiation shown in Figure S2A prompted us to investigate whether further enhancement of Wnt pathway activity during the differentiation phase could further advance TRC differentiation. Keeping CHIR99021 at 3 μM, we increased the proportion of R-spondin 1-conditioned medium (R-spondin 1 CM) in the differentiation medium from 10% to 80% (v/v), which significantly enhanced canonical Wnt pathway activation as measured by TCF/LEF reporter assay (Figure S2B). This condition was designated the 2-phase high-R-spondin method (2-phase HR; Figure 2A). Accordingly, the initially established 2-phase condition is hereafter referred to as the 2-phase low-R-spondin method (2-phase LR; Figure 3). Immunofluorescence analyses showed that the 2-phase HR condition further increased the number of POU2F3^+^, PLCβ2^+^, and GNAT3^+^ cells compared with the 2-phase LR condition (Figures 2B-D). Many of these differentiated TRCs exhibited an elongated spindle-like morphology resembling the mature TRCs in native taste buds (Figure S2C), and were positioned near the organoid surface (Video S1). In a subset of GNAT3^+^ cells, apically localized microvilli-like protrusions were also observed, suggesting morphological maturation toward functional chemosensory cells (Figure S2D). The observation that higher levels of R-spondin 1 further promoted TRC differentiation suggested that robust Wnt signaling is a critical requirement for efficient TRC differentiation.

**Figure 3.**
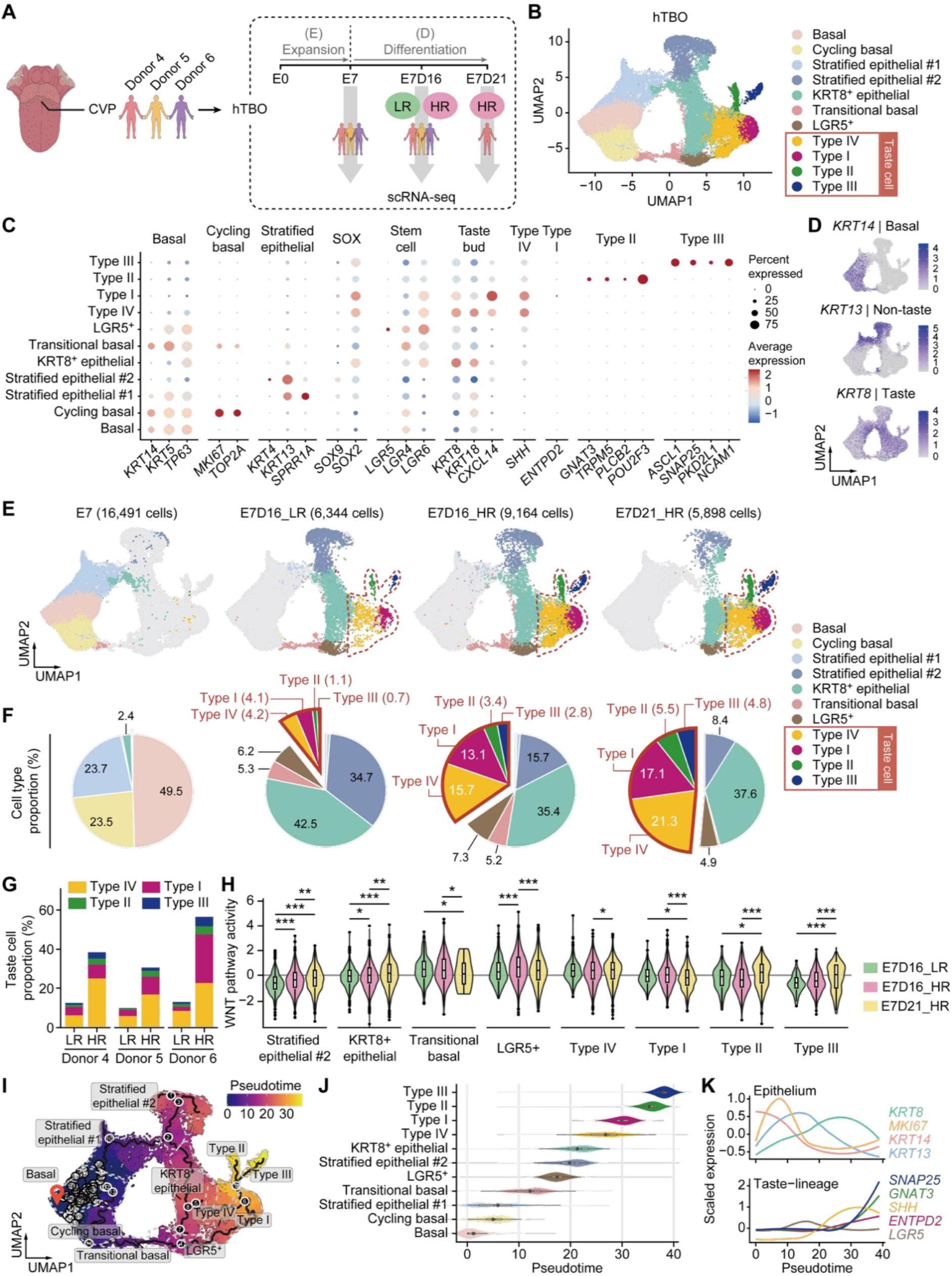
Single-cell RNA-seq analysis of hTBOs. (A) Experimental scheme for single-cell RNA-seq (scRNA-seq) of hTBOs. Organoids were generated from circumvallate papillae (CVP) of three individuals (Donors 4-6). Samples were collected at the early expansion stage (E7) and at differentiated stages under either low or high R-spondin conditions (E7D16_LR and E7D16_HR, 7 days of expansion followed by 16 days of differentiation under low or high R-spondin, respectively; E7D21_HR, 7 days of expansion followed by 21 days of differentiation under high R-spondin). (B) Integrated UMAP visualization of scRNA-seq profiles from all donors and differentiation conditions, annotated according to cell type populations. (C) Dot plot of representative marker genes used to annotate epithelial and taste-lineage cell populations (basal, cycling basal, stratified epithelial, transitional basal, KRT8^+^ epithelial, LGR5^+^, Type IV, Type I, Type II, and Type III cells). Dot size indicates the percentage of cells expressing each gene, and color indicates scaled average expression. (D) Feature plots showing expression of representative markers. *KRT14* (basal), *KRT13* (non-taste epithelium), and *KRT8* (taste differentiated). (E) UMAP visualization of samples split by differentiation stage and condition, with the number of cells in each condition indicated. (F) Pie charts depicting cell-type composition across differentiation stages and conditions. (G) Donor-wise taste cell type compositions under LR and HR conditions, shown as stacked bar plots. (H) Violin plots depicting PROGENy-inferred Wnt pathway activity scores in major epithelial and taste-lineage populations across differentiation stages and culture conditions. \**P* < 0.05, \*\**P* < 0.01 and \*\*\**P* < 0.001 by Kruskal–Wallis test followed by Dunn’s post hoc test with Bonferroni correction. (I) Pseudotime embedding of the inferred differentiation trajectory, showing progression from basal cell states through intermediate progenitor populations toward differentiated cell fates. (J) Distribution of annotated cell populations along pseudotime. (K) Expression dynamics of representative epithelium (*KRT8, MKI67, KRT14,* and *KRT13*) and taste lineage-associated genes (*SNAP25, GNAT3, SHH, ENTPD2,* and *LGR5*) plotted along pseudotime.

To determine whether the early differentiation phase exhibited greater Wnt-stimulating activity than the expansion phase, we used qRT-PCR to compare the expression of seven canonical Wnt target genes between organoids kept in expansion medium for 5 additional days after the initial 7-day expansion period (E7E5) and those switched to HR differentiation medium for 5 days (E7D5). All seven Wnt target genes, including *AXIN2, SP5, NKD1, NOTUM, RNF43, ZNRF3*, and *EPHB3*, were expressed at significantly higher levels in E7D5 organoids than in E7E5 organoids, indicating that canonical Wnt signaling was stronger in hTBOs under HR differentiation conditions than under expansion conditions (Figure S2E). Together, these findings suggest that elevated canonical Wnt activity may be required during TRC lineage specification and/or differentiation, at a level higher than that required for stem/progenitor cell expansion.

We next examined TRC differentiation dynamics by longitudinal qRT-PCR profiling of lineage-specific marker genes during the differentiation phase. The 2-phase HR condition induced a progressive and coordinated upregulation of markers associated with Type I (*ENTPD2*), Type II (*TAS1R1, TAS1R2, TAS1R3, TAS2R38, POU2F3, PLCB2,* and *TRPM5*), and Type III (*SNAP25* and *CA4*) TRCs (Figure 2E). By contrast, expression of these genes remained low or increased only modestly in 1-phase cultures (Figure 2E), consistent with the immunofluorescence data (Figures 1F and 2D). Together, these findings establish a robust 2-phase culture framework that efficiently drives TRC differentiation in hCVP-derived hTBOs, with the 2-phase HR condition producing the most pronounced enrichment of mature TRC populations.

### Single-cell RNA-seq defines stage- and condition-dependent differentiation of hTBOs

To define how cellular states evolve across the 2-phase expansion-to-differentiation process, we performed single-cell RNA sequencing (scRNA-seq) on hTBOs derived from three independent donors. Organoids were harvested at three time points across an expansion-to-differentiation time course, including the end of expansion (E7), intermediate differentiation under LR or HR conditions (E7D16_LR and E7D16_HR), and extended differentiation under HR conditions (E7D21_HR) (Figure 3A).

After quality control, we obtained 37,897 cells across all conditions (16,491 E7 cells, 6,344 E7D16_LR cells, 9,164 E7D16_HR cells, and 5,898 E7D21_HR cells). Unsupervised clustering resolved 24 transcriptionally distinct epithelial clusters (Figures S3A-S3D). Based on canonical marker gene expression, we annotated 11 epithelial populations, including basal cells, cycling basal cells, stratified epithelial cells #1 and #2, transitional basal cells, LGR5⁺ cells, KRT8⁺ epithelial cells (*KRT8⁺* but TRC-negative), and differentiated Type I-IV TRCs (Figures 3B, 3C, and S3E). Importantly, cells expressing *KRT14* (basal), *KRT13* (non-taste lingual), and *KRT8* (taste epithelial) were largely non-overlapping, suggesting that basal stem/progenitor cells may give rise to two discrete epithelial fates, non-taste lingual and gustatory epithelial states, which are maintained as distinct lineages in human taste buds, as observed in mice^22,23^ (Figure 3D). Each TRC subtype, defined by established subtype-specific markers, occupied a distinct region within the UMAP space (Figure S3E).

The relative abundance of the annotated epithelial populations changed markedly upon transition from the expansion to the differentiation phase, demonstrating that the 2-phase culture system efficiently drives organoid cells toward TRC fates (Figures 3E and 3F). E7 organoids were initially dominated by basal and non-taste lingual epithelial populations, whereas later differentiated cultures (E7D16_LR/HR, E7D21_HR) exhibited marked depletion of these populations, accompanied by a concomitant expansion of LGR5⁺ cells and TRC populations. Notably, we also identified an annotated cell population positioned between cycling basal and LGR5⁺ cells, termed “transitional basal cells”, which likely represents an intermediate state along the basal-to-LGR5⁺ lineage trajectory, capturing the transition from primitive stem/progenitor cells to more lineage-committed states.

This composition shift was most pronounced under the HR condition, which further reduced the stratified epithelial #2 and KRT8^+^ epithelial populations while increasing the differentiated TRC populations from 10.1% (E7D16_LR) to 35% (E7D16_HR) (Figure 3F). Extending the differentiation period under the HR condition from E7D16 to E7D21 led to an additional increase in TRC abundance, reaching nearly half of all cells (48.7%), accompanied by the marked depletion of the transitional basal population, suggesting continued progression from progenitor states toward mature TRC fates over time (Figure 3F). TRC enrichment under the HR condition was reproducible across all three donors (Figures 3G and S3F). Together, these data indicate that epithelial-state progression and gustatory cell production are strongly influenced by R-spondin 1 levels through enhanced responsiveness to Wnt ligands. In parallel, differentiation becomes progressively more pronounced as the culture period is extended.

To identify signaling pathways associated with TRC differentiation in hTBOs, we applied PROGENy to estimate the activity of 14 key signaling pathways across the distinct cell populations resolved by single-cell transcriptomics^43^. Notably, Wnt pathway activity was elevated in transitional basal cells, LGR5⁺ cells, and Type IV cells, which lie along the differentiation continuum from cycling basal cells to early TRCs^21,22^ (Figures S3G and S3H). At E7D16, Wnt pathway activity peaked in the LGR5⁺ cell compartment and was further enhanced under HR conditions, placing LGR5⁺ cells at the center of the epithelial response to elevated R-spondin 1 signaling (Figure 3H). However, this enhancement was attenuated at E7D21_HR, suggesting that Wnt dependence may decrease as cells approach terminal differentiation.

Finally, trajectory analysis ordered epithelial populations along pseudotime, starting from basal and cycling basal cells, progressing through transitional basal and LGR5⁺ cells, and ending in differentiated taste cell fates (Figures 3I and 3J). Along this trajectory, basal and proliferative markers (*MKI67* and *KRT14*), as well as the non-taste epithelial marker *KRT13*, were most highly expressed at early pseudotime and progressively declined thereafter. *LGR5* expression emerged at intermediate pseudotime, followed by induction of taste-lineage markers (*KRT8* and *SHH*) and, subsequently, mature TRC markers (*ENTPD2* for Type I, *GNAT3* for Type II, and *SNAP25* for Type III) at late pseudotime (Figure 3K). Collectively, these single-cell analyses support the presence of molecularly distinct stem/progenitor populations, likely comprising basal, cycling basal, transitional basal, and LGR5⁺ cells, that ultimately contribute to the generation of multiple mature TRC subtypes.

In summary, HR culture conditions enhanced TRC differentiation and increased Wnt pathway activity, particularly in transitional basal, LGR5⁺ and type IV cell populations, while producing consistent differentiation trajectories across donors. These findings establish our 2-phase culture system as an effective strategy for generating human TRCs and point to a potential link between enhanced Wnt signaling in putative stem/progenitor populations and increased TRC differentiation.

### Cross-species analysis reveals species-divergent LGR5⁺ cell identities

Mouse taste tissue has served as the primary model for understanding taste cell lineage specification and subsequent differentiation^44–46^. However, the extent to which transcriptional identities or programs are conserved across taste lineages between mouse and human remains poorly understood. To assess this, we performed cross-species integrative analysis with published mouse taste tissue^46^ and mouse taste organoid datasets^38^. We first annotated cell types in the mouse datasets based on canonical marker gene expression, using the same framework applied to hTBOs (Figures S4A–S4D). Using canonical correlation analysis (CCA)^47^, we projected hTBO cells together with mouse cells derived from taste tissue or taste organoids into a shared embedding space (Figures 4A and 4C). This analysis revealed broad conservation of lingual and gustatory epithelial states, with high cosine similarity (>0.8) among cycling basal, basal, stratified epithelial #1 and mature TRC populations, indicating conserved transcriptional identity across species (Figures 4B and 4D). Notably, however, LGR5⁺ cells showed weaker cross-species similarity when compared with both mouse taste tissue and organoid datasets, suggesting species-dependent divergence in their gene expression programs. By contrast, strong concordance was observed between the mouse taste tissue and organoid datasets across most taste lineages, including LGR5⁺ cells (Figures S4E and S4F). To define the molecular features underlying divergent LGR5⁺ cell identities, we first identified LGR5⁺ cell signature genes within each species and then compared the resulting signatures between humans and mice. For each species, LGR5⁺ cell signatures were derived by differential expression analysis comparing LGR5⁺ cells with all other cell types. Genes were retained if they met the following criteria: adjusted *P* < 0.05, log₂ fold change > 1, and expression in more than 20% of cells. Apart from *LGR5* itself, overlap between human and mouse signatures was limited, indicating that LGR5⁺ cells in the two species are characterized by distinct transcriptional programs (Figure 4E). Indeed, two signatures included distinct transcriptional regulators, such as *ERG* in human LGR5⁺ cells and *Foxe1* in mouse Lgr5⁺ cells (Figure 4F). Notably, Wnt pathway–associated genes were differentially represented between the human and mouse signatures, with hTBO LGR5⁺ cells showing stronger and more broadly distributed expression of Wnt-associated genes (Figures 4E and S4G). Although human and mouse LGR5⁺ cells differed broadly in their overall signature gene representation, this divergence was more pronounced among Wnt-associated genes, whereas other conserved functional categories, including proliferation and morphogenesis, were comparably represented between the two species (Figure S4G). These findings raise the intriguing possibility that heightened Wnt pathway activity may be a principal driver of the distinctive transcriptional program of human LGR5⁺ cells. Nevertheless, further studies with human taste bud tissue will be required to determine whether the strong Wnt stimulation used for efficient differentiation of hTBOs also contributes to the observed difference between species.

**Figure 4.**
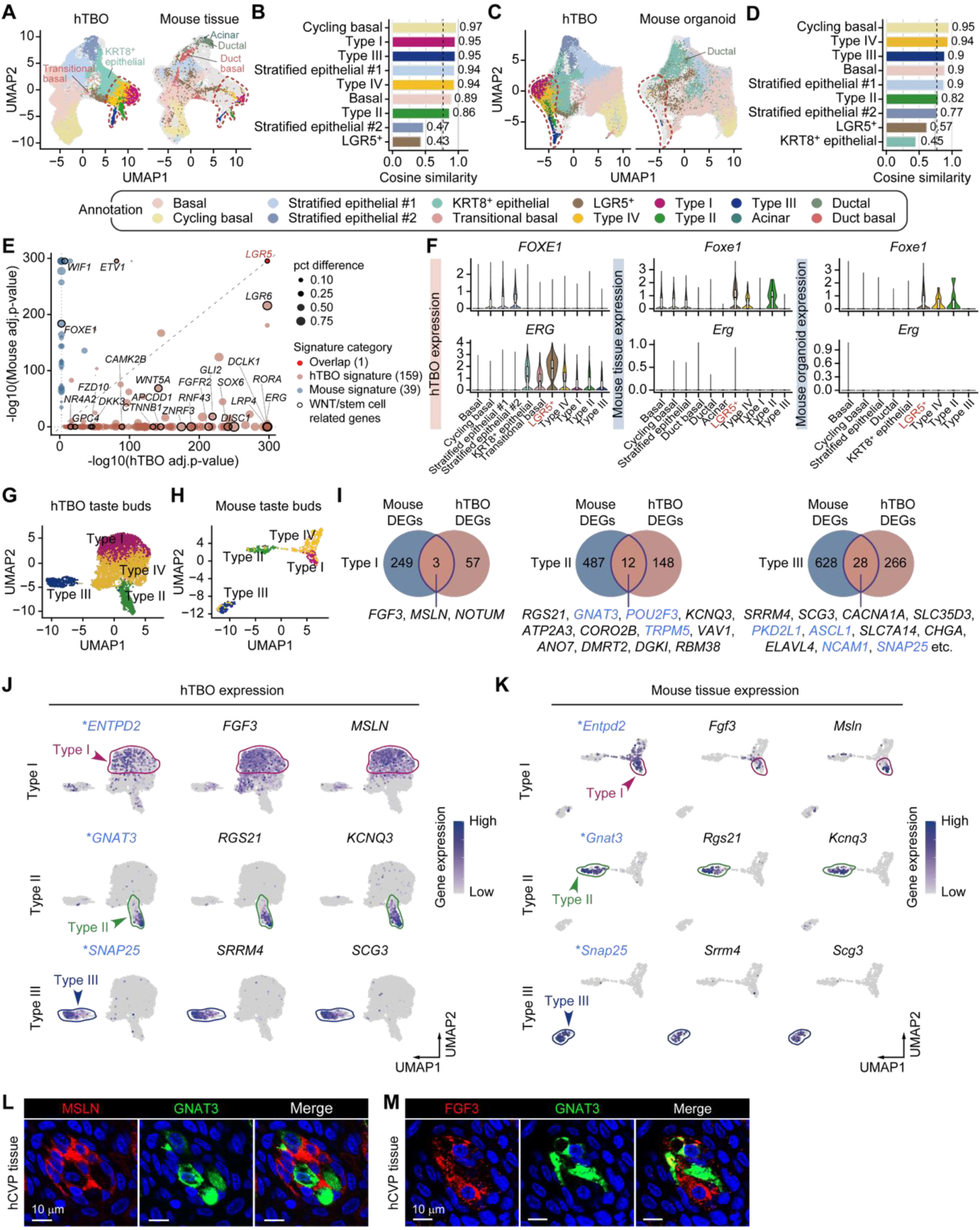
Cross-species comparison reveals conserved and subtype-specific transcriptional features of HR-derived hTBO taste cells. (A, C) UMAP embeddings generated from CCA-based integration of hTBOs with public mouse posterior tongue tissue (A) and organoid (C) datasets (GSE274014 and GSE191169, respectively), with cells colored according to annotated cell type identities. Cell types not shared between hTBOs and mouse datasets are labeled. (B, D) Cosine similarity scores calculated in CCA space between matched hTBOs and mouse taste tissue (B) or mouse organoid (D) cell populations. The dotted line indicates a cosine similarity of 0.8. (E) Scatter plot of LGR5⁺ cell signature genes identified in hTBOs and mouse taste tissue. The x- and y-axes show −log_10_ (adjusted p-value) in hTBOs and mouse tissue, respectively. Signature genes were defined as adjusted P < 0.05 (Wilcoxon rank-sum test with Benjamini-Hochberg correction) and expressed in > 20% of cells in each dataset. Dot size represents the difference in the percentage of expressing cells between the two species in LGR5^+^ cells (hTBOs - mouse tissue for hTBO signature genes; mouse tissue - hTBO for mouse tissue signature genes), with larger dots indicating stronger species-biased expression. (F) Cell type-specific expression levels of representative LGR5⁺ cell signature genes, *Foxe1* (mouse) and *ERG* (human), across hTBO, mouse tissue, and mouse organoids. (G, H) UMAP plots showing annotated taste cell subtypes in hTBO HR-derived taste buds (G) and mouse tissue taste buds (H). (I) Venn diagrams summarizing overlaps between marker genes identified in mouse taste tissue and hTBOs for Type I, Type II, and Type III taste cells. Representative shared and species-enriched genes are listed for each subtype. Blue-highlighted genes indicate human genes whose mouse orthologs are established TRC markers. (J, K) Feature plots showing the expression of representative candidate of human-mouse conserved markers in hTBO TRCs (J) and mouse tissue TRCs (K), including *ENTPD2*, *FGF3*, and *MSLN* in Type I cells; *GNAT3*, *RGS21*, and *KCNQ3* in Type II cells; and *SNAP25*, *SRRM4*, and *SCG3* in Type III cells. Asterisks indicate established marker genes for each taste cell subtype. (L) Representative immunofluorescence images show co-staining of candidate Type I TRC marker MSLN and Type II TRC marker GNAT3 in hCVP tissue. Scale bars, 10 μm. (M) Representative immunofluorescence images show co-staining of candidate Type I TRC marker FGF3 and Type II TRC marker GNAT3 in hCVP tissue. Scale bars, 10 μm.

A recent study in mice identified FOXE1 as a transcription factor enriched in posterior tongue Lgr5⁺ cells and showed that its loss impairs taste and salivary cell differentiation^46^. Consistent with this report, *Foxe1* was enriched in mouse Lgr5⁺ cells across both tissue and organoid datasets (Figure 4F). In contrast, *FOXE1* expression was absent in hTBO LGR5⁺ cells and instead detected only at low levels in cycling basal and stratified epithelial populations, suggesting that *Foxe1* may have a mouse-biased role in LGR5⁺ cells (Figure 4F). To identify transcriptional regulons of human and mouse LGR5⁺ cells, we performed gene regulatory network analysis using SCENIC^48^ (Figure S4H). Several transcription factors, including *ERG*, *RORA*, and *SOX6*, which were also included among the hTBO LGR5^+^ signature genes, were identified as significant hTBO-specific regulons of LGR5⁺ cells (Figure 4F and S4I). By contrast, in both mouse taste tissue and mTBOs, these factors showed minimal expression or were detected mainly in non-Lgr5⁺ cell populations (Figures 4F and S4I). Together, these findings indicate that human and mouse LGR5⁺ cells constitute transcriptionally distinct states, highlighting the utility of hTBOs for interrogating human-specific features of stem cell-driven taste bud regeneration and turnover.

### Cross-species comparison identified novel conserved markers of human TRC subtypes

The paucity of validated human TRC markers has limited the cellular and molecular characterization of human TRCs^49,50^. To overcome this barrier, we leveraged cross-species comparative analyses to identify novel TRC markers conserved across humans and mice. We first subset the four TRC populations (Type I, Type II, Type III, and Type IV) from the hTBO and mouse taste tissue datasets, enabling a focused analysis of subtype-specific marker expression within each population (Figures 4G and 4H).

We identified genes that were highly differentially expressed among Type I, Type II, and Type III cells in both hTBOs and mouse taste tissue, and subsequently extracted the subset shared between the two species (Figure 4I). This analysis revealed 3, 12, and 28 top DEGs shared between hTBOs and mouse taste tissue for Type I, Type II, and Type III populations, respectively (Figure 4I). Among these shared genes, the previously validated mouse marker genes *Gnat3*, *Pou2f3*, and *Trpm5* for Type II cells and *Pkd2l1*, *Ascl1*, *Ncam1*, and *Snap25* for Type III cells were recovered, providing internal validation of our cross-species approach. Notably, several shared genes exhibited stronger expression than established TRC markers, suggesting the existence of more robust novel subtype-specific markers (Figures 4J, 4K, S4J and S4K).

Among these candidate genes, we focused initially on validating Type I markers because relatively few shared genes were identified for the Type I population and, notably, *ENTPD2*, the human ortholog of the well-established mouse Type I marker *Entpd2*, did not meet the DEG selection criteria applied to hTBOs due to the low proportion of expressing cells. Of the three genes identified—*MSLN*, *FGF3*, and *NOTUM*—we performed immunostaining for MSLN and FGF3 using specific antibodies in human native taste bud tissue. We found that MSLN and FGF3 exhibited specific immunoreactivity in subsets of hCVP taste bud cells, and these signals were distinct from GNAT3 immunoreactivity observed in Type II cells (Figures 4L and 4M). Because available ENTPD2 antibodies that successfully detect the protein in mouse taste bud tissue did not produce specific staining in human taste tissue, we instead performed ENTPD2 and MSLN co-staining in mouse CVPs. MSLN immunoreactivity colocalized with that of the canonical Type I TRC marker ENTPD2 in mouse tissue, providing additional support for MSLN as a Type I TRC marker conserved between human and mouse taste tissue (Figure S4L). Together, these findings demonstrate the utility of cross-species analysis of hTBOs and mouse taste tissue for identifying novel markers of TRC subtypes and suggest that this strategy can be extended to discover additional conserved and species-specific markers for TRC subtypes beyond Type I cells.

### hTBOs generate functionally responsive TRCs

We next asked whether hTBO-derived TRCs are functionally responsive to taste stimuli. Calcium imaging of Fluo-4-loaded hTBOs differentiated under the 2-phase HR condition revealed that denatonium, a bitter tastant, elicited robust calcium transients in subsets of cells (Figure 5A). Because the three-dimensional architecture of hTBOs limited reliable Fluo-4 loading and optical detection primarily to dye-loaded cells along the rounded periphery of organoids attached to the coverslip, ROI analysis was restricted to cells in this region. Under these conditions, approximately 10–30% of dye-loaded ROIs responded to 20 mM denatonium, indicating that a subset of hTBO cells acquire functional bitter responsiveness (Figures 5A and 5B). Denatonium evoked dose-dependent calcium responses, supporting the specificity of the response (Figure 5B), and repeated denatonium stimulation produced reproducible responses (Figure 5C). These patterns were consistently observed across independent donors, indicating that bitter responsiveness we observed is not donor-specific (Figures 5C and S5A). Moreover, treatment with the PLC inhibitor U-73122 markedly attenuated denatonium-evoked calcium responses, demonstrating that bitter signaling in hTBOs depends on the canonical PLC-mediated taste transduction pathway (Figures 5D and S5B).

**Figure 5.**
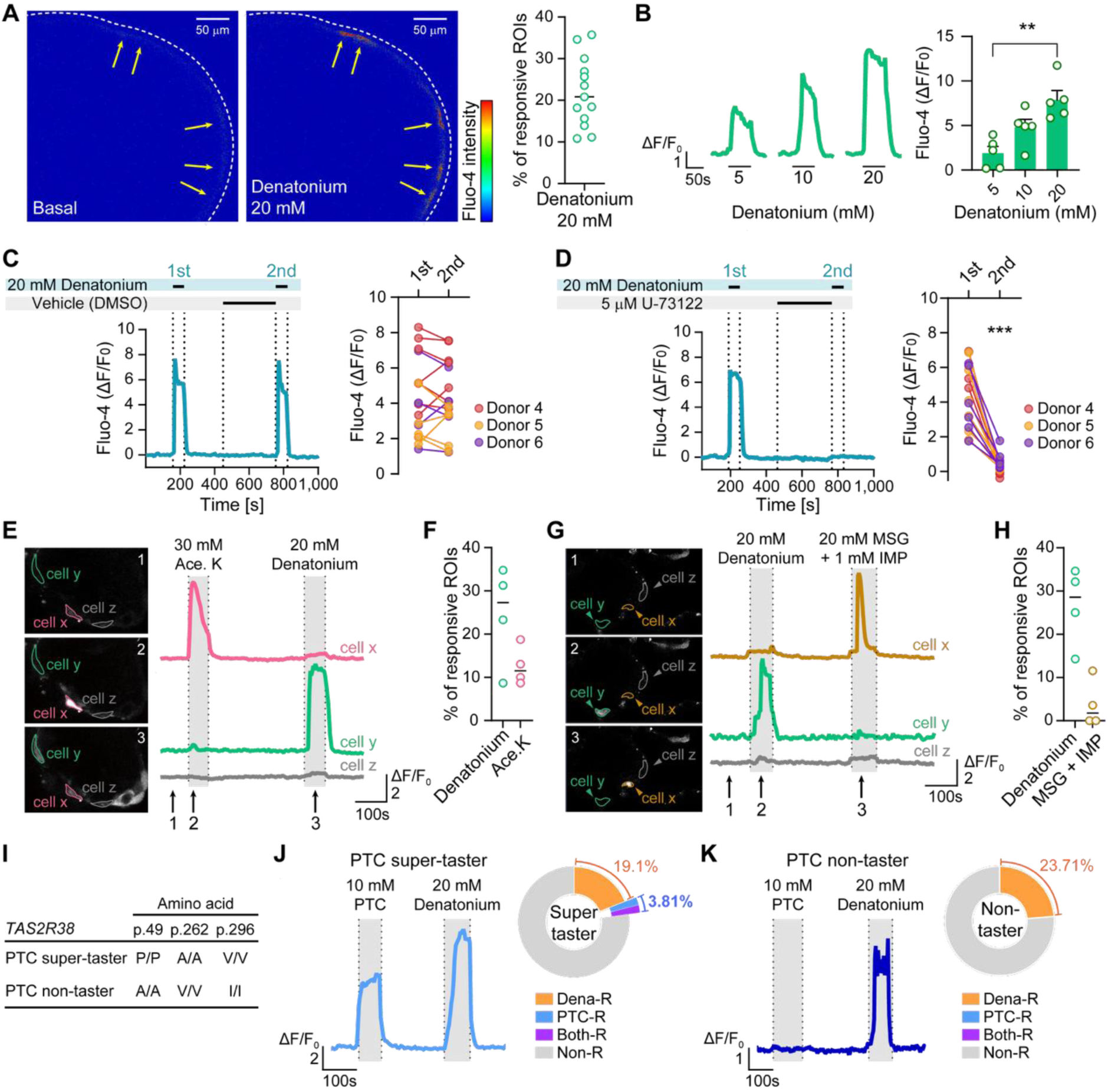
Functional characterization of taste responses in hTBOs. (A) Representative fluorescence image of Fluo-4 AM-loaded hTBOs during basal buffer perfusion and 20 mM denatonium stimulation (left). Yellow arrows indicate regions of interest (ROIs) exhibiting Ca^2+^ responses upon denatonium stimulation. The dashed line outlines a hTBO. Percentage of ROIs responsive to 20 mM denatonium (right; n = 13 independent organoid cultures from three independent donors). (B) Representative Ca^2+^ trace of Fluo-4-loaded hTBOs stimulated with increasing concentrations of denatonium (left). The solid line indicates the period of denatonium application. Quantification of peak Fluo-4 calcium responses to taste stimulation (ΔF/F_0_; right; n = 5 ROIs from a single donor). \*\**P* < 0.05 compared with 5 mM; one-way ANOVA followed by post hoc Bonferroni correction. (C, D) PLC dependence of denatonium-evoked Ca^2+^ responses in hTBOs. Representative Ca²⁺ traces of hTBOs stimulated with 20 mM denatonium after treatment with vehicle (DMSO; C) or the PLC inhibitor U-73122 (5 µM; D). Paired dot plots show peak ΔF/F_0_ values elicited by the first and second stimulations (right; n = 16 ROIs for C and n = 18 ROIs for D, from three independent donors). Colors denote organoids derived from Donors 4–6. \*\*\**P* < 0.001, paired Student’s *t* test (first vs. second stimulation). (E, G) Representative fluorescence images of Fluo-4-loaded hTBOs at the indicated time points (left) and corresponding Ca^2+^ traces for cells x, y, and z marked in the images (right). (E) Numbered arrows indicate the timing of (1) basal buffer perfusion, (2) stimulation with 30 mM acesulfame potassium (Ace.K), and (3) stimulation with 20 mM denatonium. (F) Percentage of ROIs responsive to the indicated tastants (n = 4 independent organoid cultures from two independent donors). (G) Numbered arrows indicate the timing of (1) basal buffer perfusion, (2) stimulation with 20 mM denatonium, and (3) stimulation with 20 mM monosodium glutamate (MSG) supplemented with 1 mM inosine monophosphate (IMP). (H) Percentage of ROIs responsive to the indicated tastants (n = 4 independent organoid cultures from two independent donors). (I) Amino acid polymorphisms of *TAS2R38* in PTC super-taster and non-taster donors. (J, K) Representative Ca^2+^ traces of hTBOs derived from the PTC super-taster donor (J) and the PTC non-taster donor (K) stimulated with 10 mM PTC and 20 mM denatonium. Proportions of ROIs classified as denatonium-responsive (Dena-R), PTC-responsive (PTC-R), responsive to both (Both-R), or responsive to neither (Non-R) are shown (n = 3 independent organoid cultures from two independent donors for J; n = 3 independent organoid cultures from a single donor for K). Data are presented as means ± SEM.

We next tested whether hTBOs respond to other taste modalities. Acesulfame potassium (Ace. K), an artificial sweetener, and monosodium glutamate plus inosine monophosphate (MSG+IMP), an umami stimulus, elicited calcium responses in subsets of hTBO cells that were distinct from denatonium-responsive cells (Figures 5E–H), indicating the presence of functionally separate sweet/umami- and bitter-responsive cell populations. To activate ENaC-mediated salt responses while minimizing osmotic changes associated with increased NaCl concentration, we used the amiloride-withdrawal protocol described by Nomura et al.^15^. Organoids were continuously perfused with basal buffer containing the ENaC inhibitor amiloride and then switched to an otherwise identical 140 mM NaCl buffer lacking amiloride. Amiloride withdrawal did not elicit discrete or reproducible calcium responses, whereas denatonium responses in the same organoids remained intact (Figure S5C). Thus, salt-evoked calcium responses were not detected under these assay conditions, although the absence of salt-responsive cells in hTBOs cannot be excluded. Meanwhile, citric acid elicited broad calcium responses that could not be unambiguously attributed to a discrete sour-responsive TRC population, consistent with previous observation in mTBOs^30^ (Figure S5D). Together, these results demonstrate that hTBOs generate TRCs responsive to at least three taste modalities: bitter, sweet, and umami.

### hTBOs recapitulate donor-specific *TAS2R38* genotype-phenotype relationships

*TAS2R38* is one of the best-characterized examples of genetic variation influencing human taste perception^51,52^. Its common PAV and AVI haplotypes are strongly associated with sensitivity and insensitivity, respectively, to the bitterness of PTC (Figure 5I). This well-established genotype-phenotype relationship provides a useful benchmark for assessing whether hTBOs preserve donor-specific genetic and functional properties. Sequencing identified hTBO lines carrying either the PAV (PTC-sensitive) or AVI (PTC-insensitive) *TAS2R38* haplotype (Figures 5I and S5E). In calcium-imaging assays, a subset of cells within hTBOs derived from PTC tasters carrying the PAV haplotype responded to PTC, consistent with the restricted expression pattern of *TAS2R38*^53^. Notably, a fraction of these cells also responded to denatonium, likely reflecting co-expression of TAS2R38 and other denatonium-responsive TAS2R receptors. Whereas PTC-taster hTBOs contained cells that responded exclusively to PTC as well as cells that responded to both PTC and denatonium, PTC-non-taster hTBOs lacked PTC-responsive cells (Figures 5J and 5K). By contrast, denatonium elicited comparable calcium responses in both PTC-taster and PTC-non-taster hTBOs (Figures 5J and 5K). Thus, hTBOs faithfully recapitulate the canonical TAS2R38-dependent response to PTC, demonstrating preservation of donor-specific genotype-phenotype relationships. These findings underscore the utility of hTBOs as a platform for investigating genetically driven inter-individual variation in human taste perception.

## Discussion

Studies of human taste buds have been severely constrained by the extremely limited availability of tissue, as therapeutic procedures involving the tongue that provide access to such tissue are performed only infrequently. Consequently, efforts to define the conditions required to maintain human taste bud cells *in vitro* have inevitably lagged behind those for other adult stem cell–based culture systems^54,55^. In this light, establishing an hTBO system capable of long-term expansion and mature differentiation carries particular significance, as such a system would open a critical gateway to more active and systematic investigation of human taste tissue. Our hTBO system, by fulfilling these requirements, therefore provides a firm and durable foothold for advancing human taste research. Building on this foundation, future investigations are also expected to accelerate more precise elucidation of the species-specific differences that distinguish human taste buds from those of other animals.

Over the past decade, multiple efforts have been made to establish adult stem cell–derived taste organoid systems^30–32,38,39^. Since the first report of mTBOs by Ren et al., the mouse-derived organoid system has served as a valuable platform for taste research^30^. More recently, taste organoid models have been extended to additional species, including pigs and non-human primates^31,32^. Compared with these earlier achievements, current hTBOs possess several distinct advantages. First, they enable the long-term expansion of taste stem/progenitor cells for more than 20 passages and over 50 weeks, with the potential for further extension. Given the limited availability of human taste tissue, this expandability and long-term maintainability substantially increase the utility of each donor sample and provide valuable opportunities for diverse investigations using otherwise scarce tissue. Second, our hTBOs appear to generate a significantly higher proportion of mature TRCs within the organoid structure. Although direct comparison with other taste bud organoid systems is difficult, several lines of evidence strongly support this possibility. Single-cell transcriptomic analysis revealed a distinct and well-resolved TRC populations comprising Type I–III TRC subclusters, a feature that had not been clearly demonstrated in previous taste organoid systems^31,38^. These differentiated TRC populations accounted for more than 27% of all cells in the pooled analysis of organoids from three donors, reaching 37% in donor 6. This enrichment was accompanied by a prominent Type IV progenitor population, suggesting that the expansion of Type I–III differentiated TRCs may be supported by an abundant pool of Type IV precursors. Consistent with this transcriptomic enrichment, immunofluorescence imaging of TRC markers revealed abundant TRC-positive cells, further supporting the notion that hTBOs generate mature TRCs more efficiently than previously reported taste organoid systems. More importantly, this high level of TRC differentiation was not merely achieved by identifying a single favorable condition. Rather, through a multidimensional optimization strategy involving both temporal culture design and graded Wnt signaling potentiation, we showed that TRC differentiation improves along the direction of increasing Wnt signaling. This relationship suggests that further refinement of Wnt pathway activation—particularly its duration, timing, and intensity—may further enhance mature TRC generation. Third, this hTBO system begins to reveal both conserved and human-specific features of taste bud biology across multiple dimensions, including cellular composition, intercellular organization, differentiation trajectories, and transcriptional programs. For example, hTBOs revealed a previously unrecognized basal cell population, which we termed “transitional basal cells”, positioned between cycling basal cells and evolutionarily conserved LGR5⁺ cells. The elevated Wnt pathway activity in transitional basal and LGR5⁺ cells, which lie along closely connected lineage trajectories, suggests that these populations may serve as pivotal transition points in TRC fate progression, providing a possible explanation for the strong dependence of TRC differentiation on heightened Wnt signaling.

In many adult stem cell–derived organoid cultures, withdrawal of Wnt signaling after organoid cell expansion promotes differentiation by reducing signals that maintain stem cell self-renewal^27^. Consistent with this concept, Lgr5⁺ intestinal stem cells have been shown to become committed to differentiation upon withdrawal of Wnt and R-spondin signals, indicating that these niche factors function to preserve stem cell self-renewal rather than to promote differentiation^56^. In line with this paradigm, human gastric organoids show increased differentiation into MUC5AC^+^ pit cells after four days of Wnt3a withdrawal^28^. Likewise, removal of Wnt3a and R-spondin 1, together with the addition of hepatocyte maturation factors including FGF19, BMP7, dexamethasone, and DAPT, promotes the maturation of human liver organoids into hepatocyte-like cells^29^. Human pancreatic organoids similarly undergo endocrine differentiation upon transfer to a differentiation medium lacking R-spondin 1^57^.

Conversely, although less common, Wnt signaling remains essential in certain contexts beyond stem cell maintenance, functioning as an instructive cue for lineage specification and differentiation. For example, in mouse cochlear organoids, canonical Wnt signaling is required not only for the expansion of Lgr5⁺ progenitors with sensory potential but also for their subsequent differentiation into sensory hair cells^58^. Similarly, a recently developed human intestinal organoid system demonstrated that epithelial differentiation can occur in the continued presence of Wnt signaling and that modulation of Wnt activity, together with Notch and BMP signaling, influences differentiation toward specific intestinal lineages, including Paneth, goblet, and enteroendocrine cells^59^. Together, these studies indicate that Wnt signaling can contribute directly to differentiation programs rather than acting exclusively as a self-renewal signal. In other contexts, the length of Wnt activation influences lineage outcomes. During human kidney organoid differentiation, the duration of CHIR99021-mediated Wnt activation determines intermediate mesoderm patterning and subsequent lineage specification between ureteric epithelial and nephron progenitor fates. Shorter Wnt activation favors ureteric epithelial differentiation, whereas prolonged activation promotes nephron progenitor specification, ultimately leading to distinct patterns of collecting duct and nephron formation^60^, suggesting that context-dependent Wnt signaling may shape developmental trajectories by biasing lineage decisions.

Wnt signaling can also function as an instructive cue for generating selected specialized cell types. In mouse intestinal organoids derived from Lgr5⁺ stem cells, inhibition of Wnt production with IWP-2 favored enterocyte and goblet cell differentiation, whereas sustained Wnt activation promoted Paneth cell differentiation, indicating that distinct intestinal epithelial cell types exhibit different dependencies on Wnt signaling^61^. Collectively, these observations highlight the diverse and context-dependent roles of Wnt signaling in regulating differentiation, lineage specification, and cell fate determination across organoid systems.

Against this backdrop, our hTBO model underscores the species-specific and context-dependent functions of Wnt signaling while clearly demonstrating the pivotal role of Wnt signaling strength in driving TRC differentiation. Efficient progression toward the hTBO TRC lineage required the removal of mitogenic signaling, whereas mTBOs continued to generate substantial populations of TRCs in the presence of EGF. Notably, the highest level of TRC differentiation in hTBOs was achieved only when Wnt signaling was further enhanced through CHIR99021 treatment in combination with a higher-than-conventional concentration of R-spondin 1 CM. In contrast, mTBOs can spontaneously generate substantial populations of differentiated TRCs even under standard Wnt-supportive conditions that are kept unchanged from culture initiation through differentiation^30,38,39^. Whether these conditions specific for hTBOs reflect their needs for prolonged maintenance of differentiation-competent progenitors, enhanced lineage specification, priming of transcriptional and epigenetic programs that facilitate subsequent TRC differentiation, or direct promotion of differentiation-associated signaling remains an important question for future investigation.

Whether stronger Wnt signaling can promote more extensive and mature TRC differentiation remains another important open question. In the present study, Wnt signaling was enhanced using up to 80% R-spondin 1 CM, the highest concentration achievable in our culture system, suggesting that further augmentation of Wnt activity within a CM-based format is technically constrained. Recent advances in Wnt engineering have enabled more potent and precisely defined activation of canonical Wnt signaling, including Wnt–AFM complexes, targeted R-spondin mimetics, and engineered Wnt agonists^62–64^. For example, Wnt superagonists that mimic combined Wnt and R-spondin 1 activity may achieve higher maximal efficacy in canonical Wnt activation than conventional or previously developed Wnt-activating strategies^63^, potentially generating larger and more diverse TRC populations with more mature features. Such studies may help identify strategies to further improve hTBOs as a platform for studying human TRCs.

The most prominent human-specific features in our hTBOs were observed in Type I TRCs and LGR5⁺ cells. *ENTPD2*, a canonical rodent Type I TRC marker^65,66^, was unexpectedly expressed in a small fraction (11%) of putative human Type I TRC cells, which were defined as an unsupervised cluster lacking markers of Type II, III, or IV TRCs. This finding indicates that this well-established marker is not suitable for identifying Type I TRCs in human taste tissue. Instead, analysis of DEGs shared between hTBOs and mouse taste bud tissue identified *FGF3* and *MSLN* as newly conserved Type I TRC markers, which were further validated in human taste bud tissue by immunostaining. Similar cross-species approaches may facilitate the discovery of additional species-specific and species-conserved TRC markers, thereby refining single-cell transcriptomic analysis of TRCs and enabling the development of genetic tools for TRC manipulation as well as immunological tools for TRC identification. Interestingly, human and mouse LGR5⁺ cells shared remarkably few signature genes, despite their proposed roles in taste cell turnover and regeneration. This divergence is unlikely to be attributable simply to differences between organoid and native tissue datasets, because both mouse taste tissue and mouse taste organoids exhibited strong concordance across most taste-lineage populations, except for LGR5⁺ cells. The essential role of *Foxe1* in mouse LGR5⁺ cells, together with its absence in human LGR5⁺ cells, provides a clear example of how these two species employ distinct transcriptional regulatory programs within analogous stem/progenitor cell populations. These findings underscore the value of hTBOs for studying human-specific mechanisms of taste epithelial renewal and dysgeusia. Understanding both the conserved and divergent features of mouse and human taste cells will provide important insights into how findings from mouse models can be translated to human biology.

Calcium imaging showed that hTBO-derived TRCs acquire functional taste responsiveness. Denatonium evoked reproducible, dose-dependent calcium transients that were suppressed by the PLC inhibitor U-73122, indicating activation of canonical PLC-dependent bitter taste signaling. This finding provides a representative example of taste-signaling mechanisms conserved between humans and mice. In our organoids, however, detecting calcium responses to NaCl and citric acid was more complicated than detecting responses to bitter, sweet, and umami tastants, because sour and salt taste signaling may not be reliably captured by calcium imaging alone. Given the limited understanding of how human taste buds detect salt, we used the amiloride-withdrawal method as an initial approach to stimulate salt-responsive pathways in our organoids^15^. However, at least in mice, the amiloride-sensitive low-salt pathway is prominent in the anterior tongue of mice^67^; therefore, the posterior origin of our human organoids from the CVP may have contributed to the lack of a detectable response. Moreover, in mice, both ENaC, an epithelial sodium channel, and CALHM1/3, a voltage-gated ATP-release channel complex, are key molecules required for salty taste signaling, and ENaC activation drives ATP neurotransmitter release through an entirely electrical mechanism without increasing calcium transients^15^. Thus, if human organoids possess mechanisms similar to the ENaC- and CALHM1/3-dependent pathways identified in mice, NaCl stimulation could induce ATP release without a detectable elevation in intracellular Ca²^+^. In contrast to NaCl, citric acid elicited calcium transients in most measured cells. However, this widespread activity is unlikely to represent a specific physiological sour response, because sour-responsive Type III cells are known to constitute only a subset of TRCs^10^, as also confirmed by single-cell transcriptomic analysis of our organoids. Sour taste is mediated by the OTOP1 proton channel, whose activation lowers intracellular pH^16^. This intracellular acidification can depolarize the membrane by inhibiting potassium channels^68^, thereby activating voltage-gated Ca²⁺ channels and producing a rise in intracellular Ca²⁺^16^. However, under the stimulation conditions used here, citric acid may induce calcium responses through nonspecific acidification-mediated mechanisms rather than through OTOP1-dependent signaling. Moreover, extracellular acidification can directly affect the fluorescence signals of calcium indicators^69^, further complicating the interpretation of sour-evoked Ca²⁺ responses. Thus, further optimization of stimulation paradigms and reporter strategies will be required to distinguish OTOP1-specific signaling from nonspecific acidification-induced responses.

Our organoid culture platform for calcium measurements, as well as broader optical approaches for detecting cellular signals, also represents an important area for future improvement. Because organoids have a three-dimensional architecture, limited optical penetration depth restricts calcium imaging primarily to cells located near the periphery of taste bud organoids, unless the organoids are dissociated. However, dissociation-based imaging of organoid-derived cells poses additional technical challenges, as well-differentiated cells isolated from organoids do not readily adhere to the culture surface or remain stably maintained *in vitro*. Because TRCs are not confined to the organoid periphery, restricting analyses to a subset of peripheral cells likely overlooks valuable functional information from the broader TRC population. Therefore, the development or adoption of engineered platforms— such as probe-based chip technologies capable of penetrating three-dimensional organoids, enabling more uniform monitoring of internal cells, and supporting controlled microfluidic tastant delivery^70,71^—would greatly expand the applicability of this organoid platform.

Several limitations should be noted. First, determining how closely human taste bud organoids resemble native human taste tissue would ideally be achieved through single-cell transcriptomic analysis of human taste tissue. However, such analyses were challenging because of limited tissue availability. Human CVPs collected during sleep apnea surgery were scarce, and typically only one or, at most, two CVPs could be obtained from a single donor. Each CVP contains an estimated 200–250 taste buds, with approximately 50–100 cells per bud^10,72^, yielding no more than about 25,000 taste bud cells even under ideal conditions. This limited cell number represents a major obstacle to generating comprehensive single-cell datasets from native human taste tissue, which would otherwise be highly valuable for understanding the characteristics of hTBOs. Emerging low-input single-cell RNA sequencing methods, such as SMART-seq3, and high-resolution spatial transcriptomic approaches, such as Xenium, CosMx, or Seq-Scope, may help address this limitation. Second, although hTBOs exhibited robust functional responses, our assays did not fully resolve all five canonical taste modalities, particularly salt and sour. In our experiments, human bitter, sweet, and umami responses appeared to follow signaling principles that were clearly established more than a decade ago in animal models, including mice. By contrast, the mechanisms underlying salty and sour taste remained less well defined until ENaC, CALHM1/3, and OTOP1 were identified as key receptors and channels in mice. Their relevance to human taste tissue has remained difficult to assess owing to the lack of an adequate human-cell experimental system. This represents a key area in which hTBOs can make a meaningful contribution. When combined with the hTBO platform, hTBO-compatible assays designed to measure ATP release from salt-responsive cells using the intensity-based ATP-sensing fluorescent reporter iATPSnFR, or to optimize sour-taste stimulation conditions using TRC-type-specific pH reporters, would provide previously unavailable opportunities to investigate human taste signaling. Third, hTBOs have not yet been evaluated in disease-relevant contexts, including chemotherapy, infection, and aging, in this study. Because our hTBOs comprise cells required for taste bud cell regeneration and taste-sensing processes, the introduction of disease-relevant stimuli and perturbations into this system could mimic disease-associated phenotypes and provide a valuable platform for studying disease pathogenesis. Future studies using improved functional assays, patient-derived organoids, and clinically relevant modeling approaches will be important for evaluating their utility in modeling human dysgeusia.

In summary, hTBOs provide a robust and scalable experimental platform for the long-term expansion of human taste stem/progenitor cells, their subsequent differentiation into TRCs, and the molecular and functional interrogation of human taste biology. By bridging the gap between animal-model studies and human gustatory biology, this system opens new opportunities for mechanistic studies of taste cell development, precision chemosensory research, and translational investigations of taste dysfunction.

## Methods

### Human tissue specimen

Human tongue tissues containing circumvallate papillae were surgically collected from patients undergoing sleep apnea surgery at Severance hospital, Seoul. The use of human samples in this study was approved by the Institutional Review Board of Yonsei University College of Medicine (4-2018-0281). Written informed consent was obtained from all participating patients prior to tissue collection.

### Tissue processing

Human tongue tissues were collected from patients and immediately transferred on ice in basal medium consisting of Advanced DMEM/F12 (Gibco, 12634028) supplemented with 1× GlutaMAX (Gibco, 35050061), 10 mM HEPES (Gibco, 11330057), and 1× antibiotics–antimycotics (Gibco, 15240062). Upon arrival, tissues were washed with wash medium (DMEM (Gibco, 10569044) supplemented with 1% fetal bovine serum (FBS; Gibco, 16000044) and 1× antibiotics–antimycotics) to remove debris and blood, and excess connective tissue was trimmed using scissors. Tissues were then minced into small fragments using scissors and incubated in pre-warmed dispase II solution (2 mg/mL dispase II (Gibco, 17105041) in DMEM) at 37°C for 10 minutes. Following enzymatic digestion, tissue fragments were gently pipetted using an end-cut 1000 µL pipette tip. The suspension was centrifuged at 1,300 rpm for 3 minutes, and the pellet was further incubated in 1 mL TrypLE Express (Gibco, 12605028) at 37°C for 30 minutes. After incubation, tissues were mechanically dissociated again by vigorous pipetting using an end-cut 1000 µL to generate a single-cell suspension. Wash medium was added up to a final volume of 10 mL, followed by centrifugation at 1,300 rpm for 5 minutes. The resulting tissue fragments and single cells were mixed with Matrigel (Corning, 356231), seeded onto pre-warmed 6-well plates, and allowed to solidify at 37°C for 20 minutes. Expansion medium for human taste bud organoids, described in detail in the following section, was then added and cultures were maintained for approximately 2–3 weeks. Once organoids reached an appropriate size, Matrigel and residual tissue clumps were removed. Organoids were dissociated into single cells using TrypLE Express and either cryopreserved in Recovery Cell Culture Freezing Medium (Gibco, 12648010) or passaged for further culture.

### Generation of conditioned medium

L-Wnt3a cell and HEK293T R-Spondin1-Fc cell lines were kindly provided by Hans Clevers (Hubrecht Institute, Utrecht, Netherlands). Cells were seeded into T150 flasks (Falcon, 355001) at a density of approximately 1.5 × 10⁶ cells per flask in 35 mL of culture medium (DMEM (Gibco, 10569044) supplemented with 10% FBS). Cells were passaged upon reaching approximately 80% confluency. For passaging, culture medium was removed and cells were washed once with 10 mL of DPBS. Cells were then incubated with 6 mL of TrypLE at 37 °C for 5 minutes. After detachment, cells were collected and pelleted by centrifugation at 1,300 rpm for 5 minutes. Cell pellets were resuspended in an appropriate volume of culture medium and reseeded onto 150-mm culture dishes (Corning, 353025) at a density of 1.5–2.0 × 10⁶ cells per dish in 20 mL of culture medium. Cells were allowed to adhere and grow until reaching approximately 80% confluency. Conditioning was initiated by removing the culture medium and replacing it with 20 mL of harvest medium per 150-mm dish. For Wnt3a conditioned medium, the harvest medium was identical to the culture medium. For R-spondin 1 conditioned medium, Advanced DMEM supplemented with 1× GlutaMAX, 10 mM HEPES, and 1× antibiotics–antimycotics was used as the harvest medium. Cells were incubated in harvest medium for 5 days for Wnt3a conditioned medium and for 7 days for R-spondin 1 conditioned medium. After conditioning, supernatants were collected into 50 mL conical tubes and centrifuged at 1,300 rpm for 5 minutes to remove cellular debris. Clarified supernatants were filtered using filter cups (Millipore Sigma, S2GPU11RE) and stored at 4 °C for up to 6 months. For long-term storage, conditioned media were aliquoted and stored at −20 °C. Cell stocks were cryopreserved using Recovery Cell Culture Freezing Medium.

### Organoid culture

Cells isolated from human circumvallate papilla were mixed with Matrigel (Corning, 356231) and seeded onto pre-warmed 24-well plates at a density of approximately 10,000 cells per well in 40 µL of Matrigel. Matrigel was solidified for 15 minutes at 37°C, then human taste bud organoid expansion medium was added to each well. The human taste bud organoid expansion medium was based on basal medium described above and supplemented with 2% (v/v) B-27 supplement (Gibco, 17504044), 1% (v/v) N2 (Gibco, 17502001), 1 mM N-acetylcysteine (Sigma-aldrich, A0737) and the following factors: 50% (v/v) Wnt3a conditioned medium (Homemade), 50 ng/mL EGF (Peprotech, AF-100-15), 100 ng/mL Noggin (Peprotech, 120-10C), 10% (v/v) R-spondin 1 conditioned medium (Homemade), 100 ng/mL FGF10 (Peprotech, 100-26), 500 nM A83-01 (Tocris, SML0788) and 10 µM Forskolin (Tocris, 1099). 10 µM Y-27632 (Tocris, 1254) was added to prevent anoikis following single-cell dissociation. For differentiation, organoids were cultured in differentiation medium composed of basal medium supplemented with 2% (v/v) B-27 supplement, 1% (v/v) N2, 1 mM N-acetylcysteine and the following factors: 100 ng/mL Noggin, 80% (v/v) R-spondin 1 conditioned medium, 100 ng/mL FGF10, 500 nM A83-01, 10 µM Forskolin and 3 µM CHIR99021 (Tocris, 4423). Plasmocin was included in both expansion and differentiation media at a final concentration of 12.5 µg/mL to avoid mycoplasma contamination. Organoid cultures were maintained with medium changes every 2–3 days and passaged every 2–4 weeks. For passaging, Matrigel-embedded organoids were collected, resuspended in 1 mL TrypLE Express and incubated at 37°C for 5 minutes. Organoids were dissociated into single cells by repeated pipetting using a 1000 µL pipette tip fitted with a 10 µL tip (approximately 10 strokes).

### Cell expansion rate analysis

A defined number of cells (*N*_0_) was embedded in Matrigel and cultured under expansion conditions. After 2–3 weeks of culture, organoids were dissociated into single cells, and the total number of recovered cells at each passage (*N_i_*) was determined. The expansion factor at the *i*-th passage (*F_i_*) was calculated as the ratio of the total cell number obtained after culture (*N_i_*) to the number of cells reseeded at the beginning of that passage (*N*_seed_). Following cell counting, a fixed number of cells (*N*_seed_) was reseeded for subsequent culture. This procedure was repeated at each passage, and expansion factors (*F_i_*) were calculated for all passages. The theoretical cumulative total cell number after the *i*-th passage (*N*_cum,*i*_) was estimated by multiplying the expansion factors across passages and applying this value to the initial seeding cell number (*N*_0_).

Expansion factor at each passage: 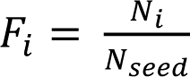

Theoretical cumulative total cell number: 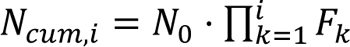

### Immunostaining

Human circumvallate papillae tissues were fixed overnight at 4 °C in 4% paraformaldehyde and subsequently cryoprotected in 30% sucrose overnight at 4 °C. Tissues were embedded in O.C.T. compound (Leica, 14020108926), frozen on dry ice, and sectioned at 10-15 µm using a cryostat (Leica). Sections were either processed immediately for immunostaining or stored at −30 °C. Prior to staining, sections were equilibrated to room temperature for 30 minutes, washed with phosphate-buffered saline (PBS) to remove residual O.C.T., and blocked with PBS containing 0.2% bovine serum albumin (BSA) and 0.3% Triton X-100 for 1 hour at room temperature. Primary antibodies were applied overnight at 4 °C, followed by three PBS washes for 10 minutes each and incubation with fluorescently labeled secondary antibodies (1:300) for 1 hour at room temperature. Nuclei were counterstained with DAPI (1:1,000) for 15 minutes at room temperature, and slides were mounted using Fluoromount-G (SouthernBiotech, 0100-01). Primary antibodies used for immunostaining are listed in Table S1.

For whole-mount immunostaining of hTBOs, organoids were washed and incubated in Cell Recovery Solution (Corning, 354253) at 4 °C for 45 minutes to remove Matrigel, then fixed in 4% paraformaldehyde at 4 °C for 45 minutes. After washing in PBT (PBS containing 0.1% Tween-20) for 15 minutes at 4 °C, organoids were blocked in organoid wash buffer (OWB; PBS containing 0.2% BSA and 0.3% Triton X-100) for 15 minutes at 4 °C. Primary antibodies diluted in OWB were incubated overnight at 4 °C with gentle shaking in 96-well plates (100 µL per well), followed by three OWB washes (200 µL per wash) and incubation with secondary antibodies (1:300 in OWB) for 1 hour at room temperature. Nuclei were counterstained with DAPI (1:1,000) for 15 minutes at room temperature. Organoids were transferred onto glass slides using an end-cut pipette tip, mounted in VECTASHIELD Antifade Mounting Medium (Vector Laboratories, H-1000-10), and coverslipped with spacers (short coverslips) placed on both sides. All images were acquired using a LSM 980 confocal microscope (ZEISS) as Z-stacks merged into three-dimensional projections.

### RNA isolation and reverse transcription quantitative PCR (qRT-PCR)

Total RNA was isolated from human taste bud organoids using TRIzol reagent (Thermo Fisher Scientific, 15596018) according to the manufacturer’s instructions. Reverse transcription was performed using a Primescript™ 1st strand cDNA synthesis kit (TAKARA Bio, 6110B) following the manufacturer’s protocol. Quantitative real-time PCR was performed on a QuantStudio™ 3 Real-Time PCR System (Applied Biosystems) using either TaqMan™ Fast Advanced Master Mix (Applied Biosystems, 4444557) or RealHelix™ Premier qPCR Kit (NanoHelix, PQH-S500). Gene expression was analyzed using TaqMan assays listed in Table S2 and SYBR Green primer pairs listed in Table S3.

### Generation of single-cell RNA sequencing data

Human taste bud organoids (hTBOs) were generated from three independent human circumvallate papillae (CVP; donors 4, 5, and 6). Organoids were collected at the early stage (E7) and at differentiated stages after culture under low- or high-R-spondin conditions (E7D16_LR, E7D16_HR, and E7D21_HR). Approximately 10⁶–10⁷ dissociated single cells from human taste bud organoids at differentiation days 0, 16, and 21 were collected for single-cell RNA sequencing. Cells were pelleted by centrifugation at 1,300 rpm for 5 minutes, resuspended in 2 mL of wash medium, divided into two LoBind tubes (Eppendorf, 0030108116), and centrifuged again at 500 × g for 5 minutes at 4 °C using a swing-bucket centrifuge. Pelleted single cells were immediately processed and fixed using the Evercode Cell Fixation Kit (Parse Biosciences, ECWT3300). Following barcode recovery, cDNA was purified and amplified by PCR according to the Evercode WT v3 platform protocol. cDNA yield and quality were assessed at multiple steps using a 4200 TapeStation (Agilent). Paired-end sequencing (PE150) was performed on a NovaSeq X Plus system (Novogene).

### Pre-processing of single-cell RNA sequencing data

Raw sequencing reads were processed using the Parse Biosciences splitpipe pipeline (v1.6.2) with the human reference genome (GRCh38) to perform read demultiplexing, barcode error correction, alignment, and gene-level quantification, generating per-sample raw count matrices. Raw count matrices were converted into Seurat objects and processed using Seurat (v5.2.0)^73^. Quality control was applied to each sample individually to exclude low-quality cells, empty droplets, and multiplets based on nCount_RNA thresholds, mitochondrial transcript proportion, and predicted doublet scores. Data were normalized using log-normalization with a scale factor of 10,000. Dimensionality reduction was performed using principal component analysis (PCA, npcs = 50). For CVP tissue samples, Harmony integration^74^ was applied across donors to correct for inter-donor batch effects. Finally, UMAP embedding was computed on the Harmony-corrected dimensional space (dims = 1:15), and cell-cell distances were calculated using FindNeighbors (dims = 1:15). Unsupervised clustering was performed using the Louvain algorithm (FindClusters, resolution = 0.3). Clusters exhibiting co-expression of immune cell and taste cell markers, indicative of potential doublets or ambiguous cell states, were excluded from downstream analyses.

### Cell-type annotation of hTBO

Clusters were annotated based on established marker genes^45,46^. Basal cells were identified by expression of *KRT14, KRT5*, and *TP63*, cycling basal cells by *MKI67* and *TOP2A*, stratified epithelial cells by *KRT4*, *KRT13*, and *SPRR1A*, LGR5^+^ cells by *LGR5*, *LGR4*, and *LGR6*, and mature taste cells by *KRT8*, *KRT18*, and *CXCL14*. The KRT8^+^ population, representing intragemmal taste cells, was further sub-clustered into distinct taste cell subtypes, including Type IV cells (*SHH*), Type I cells (*ENTPD2*), Type II cells (*GNAT3*, *TRPM5*, *PLCB2*, and *POU2F3*), and Type III cells (*ASCL1*, *SNAP25*, *PKD2L1*, and *NCAM1*).

### Pathway activity and trajectory analysis

Pathway activity scores were estimated using PROGENy^43^ and compared across annotated epithelial and taste cell populations. Differences among cell types were evaluated using the Kruskal–Wallis test followed by Dunn’s post hoc test with Bonferroni correction for multiple comparisons. Pseudotime trajectory analysis was performed using Monocle3 on the UMAP embedding derived from the Harmony-corrected PCA space, with the root node assigned to the basal cell cluster based on prior biological knowledge^75^.

### Integration with mouse tissues and organoids

To compare human and mouse taste cell transcriptomes, publicly available mouse scRNA-seq datasets were obtained, including posterior papilla tissue data (GSE274014; Verweij et al., 2025^46^) and posterior taste organoid data (GSE191169; Adpaikar et al., 2022^38^). Mouse datasets were annotated using orthologous marker genes corresponding to those applied to hTBOs, supplemented with salivary cell markers (*Dcpp1* and *Cldn2*) reported in the original study.

To identify conserved and species-specific transcriptomic features, human and mouse gene expression matrices were matched on one-to-one orthologous genes identified using biomaRt and CCA integration^76^ was performed using Seurat (dims = 1:20). Transcriptional similarity between matched human and mouse cell populations was quantified by cosine similarity in the CCA space. Differentially expressed genes for each cell population were identified using FindAllMarkers with a Wilcoxon rank-sum test, selecting genes with adjusted P < 0.05, log_2_ fold change > 1, and pct. > 0.2 as the signature within each species. Shared signatures between hTBO and mouse were designated as conserved, whereas those detected only in the hTBO or mouse dataset were classified as hTBO-enriched or mouse-enriched, respectively. To further validate the transcriptional differences at the gene regulatory network level, pySCENIC^48^ was applied to infer transcription factor regulon activity across LGR5^+^ cells from both species, identifying species-specific regulatory programs consistent with the observed signature gene expression.

### Ca^2+^ assay using Fluo-4 dye and perfusion

Twelve-millimeter coverslips were flame-sterilized and placed into 24 well plates. Coverslips were coated with 500 µL of a mixture containing 1 mg/mL poly-D-lysine (Thermo Scientific, A3890401) and 5 µg/mL laminin (Thermo Scientific, 23017015) and incubated at room temperature for 2 hours. After incubation, the coating solution was removed and the coverslips were air-dried. hTBOs were washed to remove Matrigel, resuspended in a small volume of organoid culture medium and plated onto the coated coverslips as small droplets (5-10 organoids per coverslip). Organoids were allowed to attach to the coverslips by incubating at room temperature for 20 minutes. To minimize organoid detachment during subsequent perfusion, culture medium was slowly added to a final volume of 400 µL per each well. Cultures were incubated overnight at 37 °C and Ca²⁺ imaging experiments were performed the following day.

Prior to Ca²⁺ imaging, the perfusion device was rinsed twice with double-distilled water. Physiological saline solution (PSS) was freshly prepared on the day of the experiment. Organoid-attached coverslips were gently washed with PSS supplemented with 1% BSA. Cells were then loaded with the calcium indicator Fluo-4 AM (Thermo Scientific, F14201) at a final concentration of 2 µg/mL in PSS for 1 hour at room temperature. The composition of PSS was as follows: 140 mM NaCl, 5 mM KCl, 1.8 mM CaCl₂, 1.4 mM MgCl₂, 11.5 mM glucose, 10 mM HEPES, 1.2 mM NaH₂PO₄, and 5 mM NaHCO₃ (pH 7.4). After dye loading, coverslips were mounted onto a perfusion chamber and continuously perfused with PSS during imaging. Ca^2+^ imaging was performed using an LSM 700 confocal microscope (ZEISS). Fluo-4 fluorescence was excited at 488 nm, and emitted fluorescence was collected using ZEN imaging software (ZEISS). Cells were assayed for their responses to tastants including denatonium benzoate (5, 10, and 20 mM), phenylthiocarbamide (10 mM), acesulfame potassium (30 mM), monosodium glutamate (20 mM) supplemented with 1 mM inosine monophosphate (IMP) and citric acid (20 mM, pH 4.0). In some experiments, 5 µM U73122 (Sigma-Aldrich, U6756) was treated for 5 minutes to assess the involvement of the PLCβ2 signaling pathway in Ca²⁺ responses. For salt taste stimulation, organoids were continuously perfused with a 140 mM NaCl-containing PSS supplemented with 1 µM amiloride and then switched to an otherwise identical buffer lacking amiloride.

## Acknowledgments

This work was supported by the National Research Foundation of Korea(NRF) grant funded by the Korea government (MSIT) (RS-2024-00406281 to C.H.K and S.J.M; RS-2024-00339611 to C.H.K.) and the InnoCORE program of the Ministry of Science and ICT (26-InnoCORE-02 to C.H.K. and S.J.M.) and the Korea Health Technology R&D Project through the Korea Health Industry Development Institute (KHIDI), funded by the Ministry of Health & Welfare, Republic of Korea (RS-2025-25459824 to H.C) and the Advanced Technology Hub Group (Grant No. D-2025-0021 to J.B.), Gangnam Severance Hospital, Yonsei University College of Medicine.

## Author Contributions

J.C. performed most of the experiments and analyzed the data. S.S.K. established the initial hTBO culture system. J.K. and H.M. performed single-cell transcriptomic analyses. V.Q.D. provided guidance for calcium imaging experiments. S.Z. contributed to manuscript writing and scientific discussion. H.C. provided human tissue samples and project guidance. J.B. supervised single-cell analysis and contributed to manuscript preparation. S.J.M. and C.H.K. conceived and supervised the project and wrote the manuscript.

