## Supplemental Data for "Generation of Human Taste Bud Organoids as a Human-Mimetic Platform for Modeling Taste Perception"

This Supplementary Information PDF contains the following:

Supplemental Figures S1-S5

Supplemental Table S1-S3

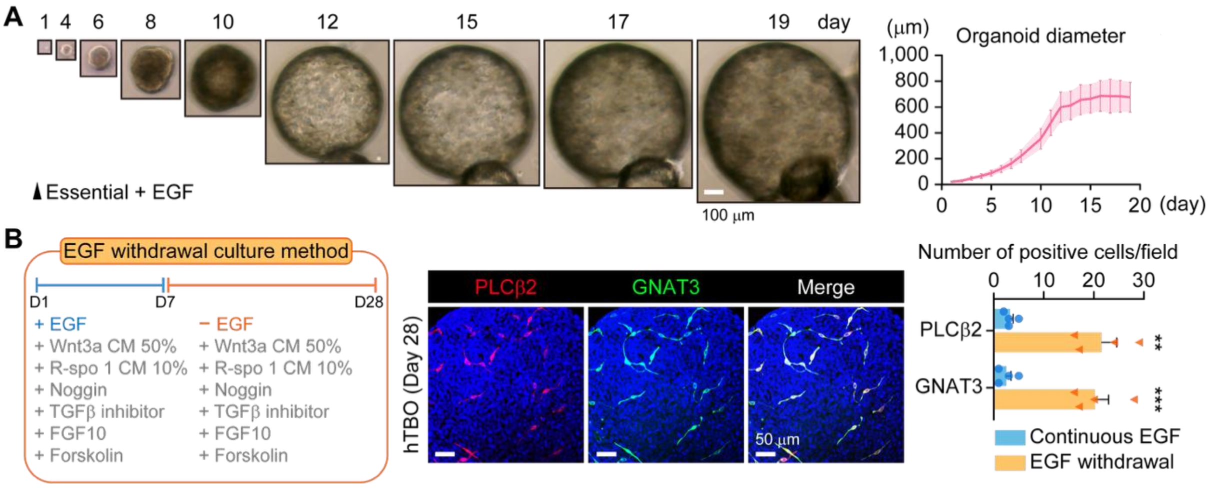


**Figure S1. Long-term expansion of hTBOs in EGF-supplemented essential medium and enhanced Type II TRC differentiation following EGF withdrawal, related to Figure 1 and Figure 2.** (A) Representative bright-field images of hTBOs cultured in essential medium supplemented with EGF at the indicated time points (left) and quantification of organoid diameter over time (right, n = 7 independent organoids from a single donor). Scale bar, 100 μm. (B) Schematic of the EGF withdrawal experiment (left). EGF was removed from the expansion medium on day 7, and cultures were subsequently maintained under EGF-withdrawal conditions until day 28. Representative immunofluorescence images of day 28 hTBOs stained for the Type II TRC markers PLCβ2 and GNAT3 (middle). Nuclei were counterstained with DAPI. Scale bars, 50 μm. Quantification of PLCβ2$^+$ and GNAT3$^+$ cells per field of view under the indicated culture conditions (right, n = 4 independent organoids from 2 independent donors). Each field of view measured 300 μm × 300 μm. $^{**}P < 0.01$ and $^{***}P < 0.001$ compared to continuous EGF; two-tailed unpaired Student's $t$ test. Data are presented as means ± SEM.

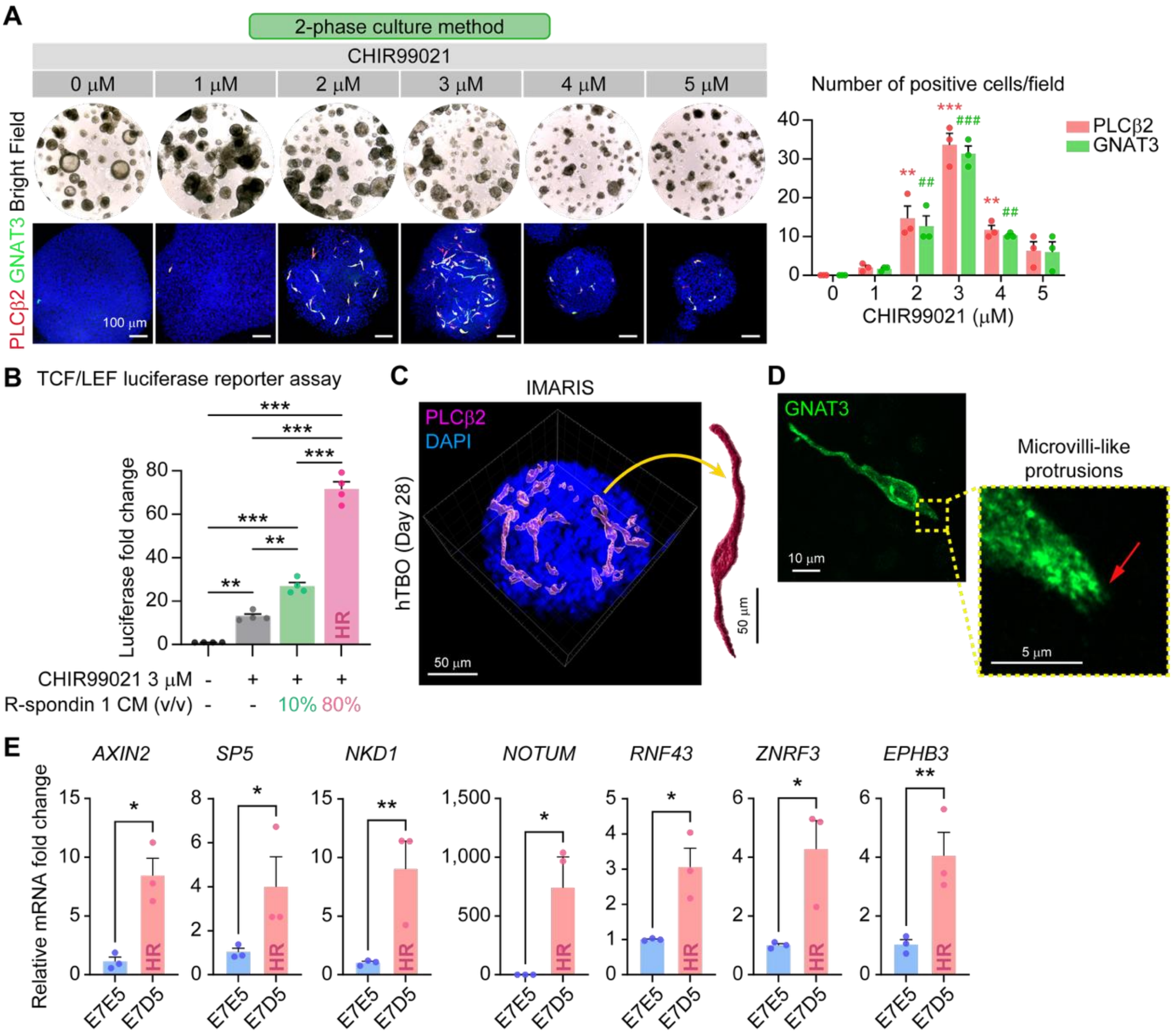


**Figure S2. Titration of CHIR99021 and R-spondin 1 defines Wnt activity levels that optimize TRC differentiation, related to Figure 2.** (A) Dose-dependent effects of CHIR99021 on hTBOs. Organoids were expanded for 7 days and then switched to differentiation medium containing 10% R-spondin 1 CM and the indicated concentrations of CHIR99021 (0–5 μM) until day 28. Representative bright-field images (top) and immunofluorescence images of individual organoids stained for PLCβ2 and GNAT3 (bottom) are shown on the left. Nuclei were counterstained with DAPI. Scale bars, 100 μm. Quantification of PLCβ2$^{+}$ and GNAT3$^{+}$ cells per field of view at each CHIR99021 concentration is shown on the right (n = 3 independent organoids from a single donor). Each field of view measured 300 μm × 300 μm. $^{**}P < 0.01$ and $^{***}P < 0.001$ for PLCβ2 compared with 0 µM, and $^{\#\#}P < 0.01$ and $^{\#\#\#}P < 0.001$ for GNAT3 compared with 0 µM; one-way ANOVA followed by post-hoc Bonferroni corrections. (B) TCF/LEF luciferase reporter assay of Wnt pathway activity under the indicated differentiation medium conditions (n = 4). The leftmost condition consisted of differentiation medium lacking both CHIR99021 and R-spondin 1 CM, and luciferase activity is expressed as fold change relative to this condition. HR, high R-spondin. $^{**}P < 0.01$ and $^{***}P < 0.001$; one-way ANOVA followed by post-hoc Bonferroni corrections. (C) Three-dimensional Imaris rendering of a day 28 hTBO generated using the 2-phase HR protocol and immunostained for PLCβ2 with DAPI counterstaining. The yellow arrow indicates an enlarged view of an individual PLCβ2$^{+}$ cell. (D) Representative immunofluorescence image of a GNAT3$^{+}$ taste receptor cell in a day 28 hTBO generated using the 2-phase HR protocol. The boxed region is shown at higher magnification. Red arrows indicate apical protrusions with microvillus-like morphology. (E) qRT-PCR analysis of Wnt target gene expression in hTBOs cultured for 7 days in expansion medium and then cultured for an additional 5 days either in expansion medium of the same composition (E7E5) or in HR differentiation medium (E7D5) (n = 3 independent organoids from 3 independent donors). $^{*}P < 0.05$ and $^{**}P < 0.01$ compared to E7E5; two-tailed paired Student's *t* test on log-transformed ratios. Data are presented as means ± SEM.

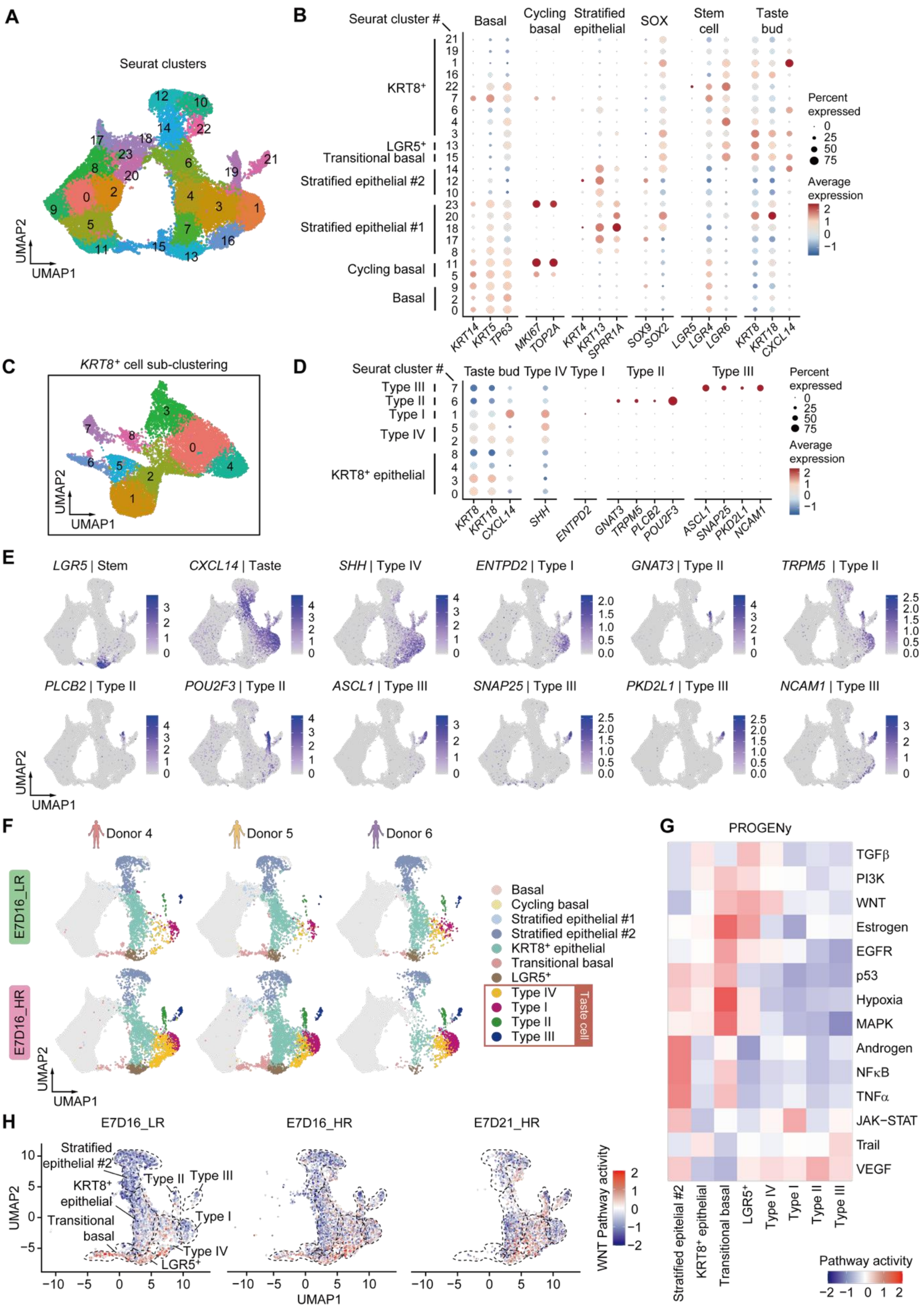
A
Seurat clusters
UMAP2
UMAP1
B
Seurat cluster #
Basal
Cycling basal
Stratified epithelial
SOX
Stem cell
Taste bud
KRT8⁺
LGR5⁺
Transitional basal
Stratified epithelial #2
Stratified epithelial #1
Cycling basal
Basal
Percent expressed
Average expression
KRT14 KRT5 TP63 MKI67 TOP2A KRT4 KRT13 SPRR1A SOX9 SOX2 LGR5 LGR4 LGR6 KRT8 KRT18 CXCL14
C
KRT8⁺ cell sub-clustering
D
Taste bud Type IV Type I Type II Type III
Type III
Type II
Type I
Type IV
KRT8⁺ epithelial
KRT8 KRT18 CXCL14 SHH ENTPD2 GNAT3 TRPM5 PLCB2 POU2F3 ASCL1 SNAP25 PKD2L1 NCAM1
E
LGR5 | Stem
CXCL14 | Taste
SHH | Type IV
ENTPD2 | Type I
GNAT3 | Type II
TRPM5 | Type II
PLCB2 | Type II
POU2F3 | Type II
ASCL1 | Type III
SNAP25 | Type III
PKD2L1 | Type III
NCAM1 | Type III
F
Donor 4
Donor 5
Donor 6
E7D16_LR
E7D16_HR
Basal
Cycling basal
Stratified epithelial #1
Stratified epithelial #2
KRT8⁺ epithelial
Transitional basal
LGR5⁺
Type IV
Type I
Type II
Type III
Taste cell
G
PROGENy
TGFβ
PI3K
WNT
Estrogen
EGFR
p53
Hypoxia
MAPK
Androgen
NFκB
TNFα
JAK−STAT
Trail
VEGF
Stratified epitelial #2
KRT8⁺ epithelial
Transitional basal
LGR5⁺
Type IV
Type I
Type II
Type III
Pathway activity
H
E7D16_LR
E7D16_HR
E7D21_HR
Stratified epithelial #2
KRT8⁺ epithelial
Transitional basal
LGR5⁺
Type II
Type III
Type I
Type IV
WNT Pathway activity
UMAP2
UMAP1

**Figure S3. Marker expression, donor-level reproducibility, and pathway activity landscape of differentiating hTBOs, related to Figure 3.** (A) UMAP visualization of scRNA-seq profiles colored by 24 Seurat clusters identified through unsupervised clustering. (B) Dot plot of representative marker genes across individual Seurat clusters used to annotate the 24 clusters into seven major cell populations (Basal, Cycling basal, Stratified epithelial #1, Stratified epithelial #2, Transitional basal, LGR5$^+$, and KRT8$^+$ cells). Dot size indicates the percentage of cells expressing each gene, and color indicates scaled average expression. (C) UMAP visualization of the KRT8$^+$ cell population from (A), colored by 9 Seurat clusters identified through further unsupervised clustering. (D) Dot plot of representative marker genes across individual Seurat clusters used to annotate the KRT8$^+$ cell subtypes (KRT8$^+$ epithelial, Type IV, Type I, Type II, and Type III). Dot size indicates the percentage of cells expressing each gene, and color indicates scaled average expression. (E) Feature plots showing expression of representative markers used to annotate taste-lineage cell populations in the integrated dataset: *LGR5* (LGR5$^+$ cell), *CXCL14* (mature TRCs), *ENTPD2* (Type I), *GNAT3*, *TRPM5*, *PLCB2*, and *POU2F3* (Type II), *ASCL1*, *SNAP25*, *PKD2L1*, and *NCAM1* (Type III), and *SHH* (Type IV). Color indicates log-normalized expression. Note that the scale is independent for each gene. (F) UMAP projections of hTBOs derived from individual donors (Donors 4-6) under low R-spondin (LR, top) and high R-spondin (HR, bottom) differentiation conditions at E7D16, colored by annotated cell population. (G) Heatmap of average activity scores for 14 PROGENy pathways across cell populations in differentiated organoids (E7D16_LR, E7D16_HR, and E7D21_HR). Basal, cycling basal, and stratified epithelial #1 populations were excluded because of insufficient cell numbers in the differentiated samples. Cell populations are ordered according to the inferred differentiation trajectory. Scores were z-scaled across populations. (H) UMAP feature plots showing PROGENy-inferred WNT pathway activity under E7D16_LR, E7D16_HR, and E7D21_HR conditions. Dashed outlines demarcate annotated cell populations, which are labeled in the leftmost panel.

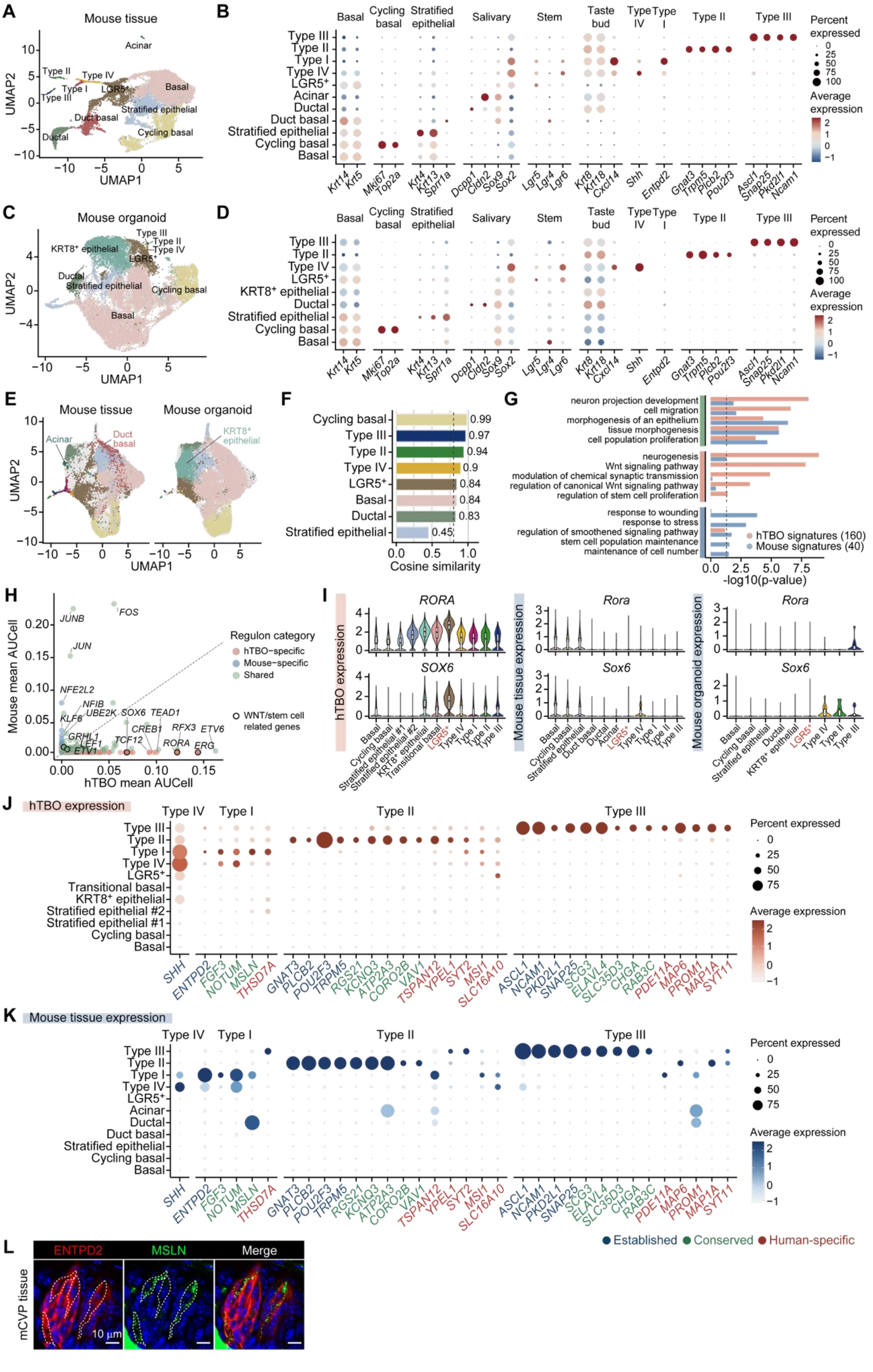
A
Mouse tissue
UMAP1
UMAP2
Acinar
Type II
Type IV
Type I
Type III
LGR5+
Basal
Stratified epithelial
Duct basal
Ductal
Cycling basal
B
Basal
Cycling basal
Stratified epithelial
Salivary
Stem
Taste bud
Type IV
Type I
Type II
Type III
Percent expressed
Average expression
C
Mouse organoid
KRT8+ epithelial
D
E
Mouse tissue
Mouse organoid
Acinar
Duct basal
KRT8+ epithelial
F
Cycling basal 0.99
Type III 0.97
Type II 0.94
Type IV 0.9
LGR5+ 0.84
Basal 0.84
Ductal 0.83
Stratified epithelial 0.45
Cosine similarity
G
neuron projection development
cell migration
morphogenesis of an epithelium
tissue morphogenesis
cell population proliferation
neurogenesis
Wnt signaling pathway
modulation of chemical synaptic transmission
regulation of canonical Wnt signaling pathway
regulation of stem cell proliferation
response to wounding
response to stress
regulation of smoothened signaling pathway
stem cell population maintenance
maintenance of cell number
hTBO signatures (160)
Mouse signatures (40)
-log10(p-value)
H
Mouse mean AUCell
hTBO mean AUCell
Regulon category
hTBO−specific
Mouse−specific
Shared
WNT/stem cell related genes
I
RORA
SOX6
hTBO expression
Rora
Sox6
Mouse tissue expression
Mouse organoid expression
J
hTBO expression
Transitional basal
Stratified epithelial #2
Stratified epithelial #1
K
Mouse tissue expression
Established
Conserved
Human-specific
L
ENTPD2
MSLN
Merge
mCVP tissue
10 μm

**Figure S4. Cross-species marker and regulon analysis of hTBOs, related to Figure 4.** (A, C) UMAP visualizations of mouse CVP dataset (GSE274014; Verweij et al., 2025[1]) (A) and the mouse CVP organoid dataset (GSE191169; Adpaikar et al., 2022[2]) (C), colored by annotated cell populations. (B, D) Dot plots of the marker genes used for cell type annotation in mouse tissue (B) and organoid (D) datasets, grouped according to the populations they define. Dot size indicates the percentage of cells expressing each gene and color indicates scaled average expression. (E) UMAP of the CCA-integrated embedding of mouse tissue and mouse organoid datasets, illustrating transcriptional correspondence between *in vivo* tissue-derived and organoid-derived cell populations. Cell types unique to either dataset are labeled to highlight species-specific populations. (F) Cosine similarity analysis in CCA space between matched cell populations from mouse tissue and organoid datasets. (G) GO Biological Process enrichment of hTBO-specific and mouse-specific $LGR5^+$ cell gene signatures. The x-axis shows $-\log_{10}$(p-value) for enriched terms. Bar color indicates the signature group: hTBO-specific (red), mouse-specific (blue). Left annotation indicates whether each term is conserved (green), hTBO-enriched (pink), or mouse-enriched (blue). The dashed line denotes the significance threshold at $-\log_{10}$(p-value) = 1. (H) Scatter plot comparing the mean AUCell scores of $LGR5^+$ cell regulons identified by SCENIC analysis in hTBO (x-axis) and mouse taste tissue (y-axis). Each dot represents a regulon and is colored according to its classification as hTBO-specific (pink), mouse-specific (blue), or shared (green). The diagonal line ($y = x$) indicates equal regulon activity in hTBOs and mouse taste tissue. (I) Cell type-specific expression of representative human $LGR5^+$ cell regulon transcription factors (*RORA* and *SOX6*) and their mouse orthologs (*Rora* and *Sox6*) across hTBOs, mouse taste tissue, and mouse organoids. (J) Dot plot summarizing expression of established and candidate taste cell markers, including conserved and human-enriched genes, in hTBOs across Type I, Type II, Type III, and Type IV populations. (K) Dot plot showing expression of corresponding established and candidate marker genes in mouse tissue-derived populations, enabling comparison of conserved versus human-enriched transcriptional features. Dot size indicates the percentage of cells expressing each gene, and color intensity reflects scaled average expression. (L) Representative immunofluorescence images showing co-localization of the Type I TRC marker ENTPD2 and the candidate Type I-associated marker MSLN in mouse CVP tissue. Scale bars, 10 μm.

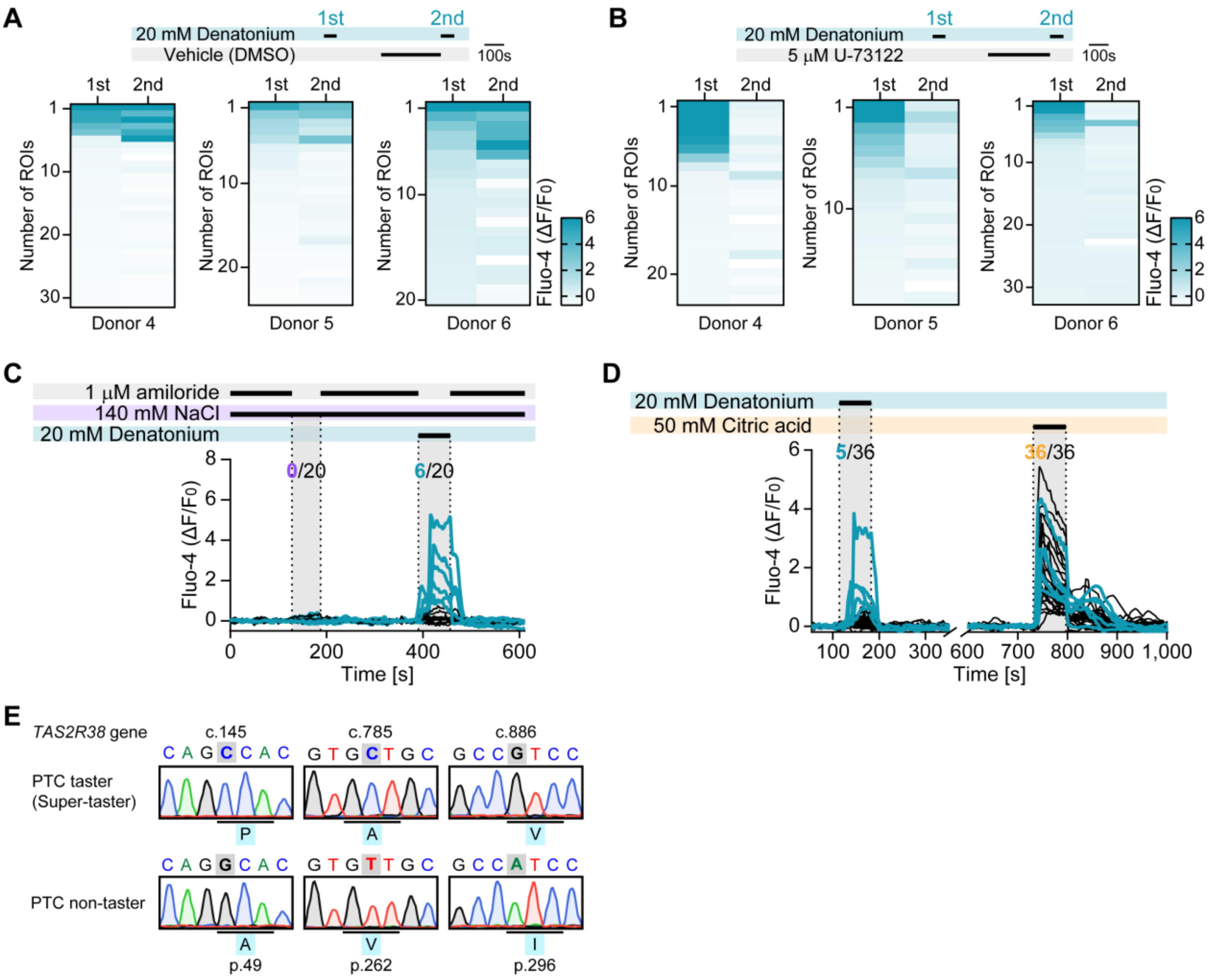


**Figure S5. Donor-level PLC dependence of bitter responses, salt and sour stimulation, and *TAS2R38* genotyping, related to Figure 5.** (A-B) Heatmaps of Fluo-4 AM $\Delta F/F_0$ responses from individual regions of interest (ROIs) in 2-phase HR-differentiated hTBOs derived from three donors (Donors 4-6). Organoids were stimulated twice with 20 mM denatonium benzoate. Between stimulations, vehicle (DMSO; A) or the PLC inhibitor U-73122 (5 µM; B) was applied for 5 min. Each row represents a single ROI, ordered according to the amplitude of the first denatonium response. Color indicates $\Delta F/F_0$. (C) $Ca^{2+}$ responses to NaCl stimulation. Amiloride (1 µM) was included in the basal buffer containing 140 mM NaCl to suppress tonic ENaC activity. NaCl stimulation was initiated by removing amiloride from the buffer. After a wash period, 20 mM denatonium benzoate was applied as a positive control. Denatonium evoked responses in 6 of 20 ROIs, whereas no ROI responded to NaCl (n = 20

ROIs from a single donor). (D) $Ca^{2+}$ responses to citric acid stimulation. A 20 mM denatonium benzoate stimulus was first applied as a positive control, followed by a wash period and stimulation with 50 mM citric acid (pH 4). Note the break in the time axis. Denatonium elicited responses in 5 of 36 ROIs, whereas citric acid elicited responses in all 36 ROIs (n = 36 ROIs from a single donor). (E) Sanger sequencing chromatograms of genomic DNA spanning the three common coding polymorphisms of *TAS2R38* (c.145G>C, p.A49P; c.785T>C, p.V262A; c.886A>G, p.I296V) in the PTC super-taster donor (PAV/PAV) and the PTC non-taster donor (AVI/AVI).

Table S1. All antibodies used for immunostaining in this study, related to STAR methods.

| **Protein name** | **Vendor** | **Catalog #** | **Host** | **Dilution ratio** |
|---|---|---|---|---|
| **KRT14** | Abcam | ab7800 | mouse | 1:200 |
| **KRT8** | Abcam | ab53280 | rabbit | 1:200 |
| **KRT5** | BioLegend | 905904 | chicken | 1:200 |
| **SOX2** | Abcam | ab92494 | rabbit | 1:200 |
| **Ki67** | CST | 9449 | mouse | 1:100 |
| **PLCB2** | Abcam | ab220285 | rabbit | 1:200 |
| **GNAT3** | Novus | NBP1-20926 | goat | 1:200 |
| **POU2F3** | CST | 36135 | rabbit | 1:200 |
| **MSLN** | Proteintech | 83744-2-RR | rabbit | 1:100 |
| **MSLN** | Neobiotechnologies | 10232-MSM1-P0 | mouse | 1:100 |
| **FGF3** | Proteintech | 16874-1-AP | rabbit | 1:100 |
| **ENTPD2** | Ectonucleotidases-ab | mN2-36LI6 | rabbit | 1:100 |

Table S2. TaqMan Assays used for qPCR, related to Figure 2.

| Gene name | Assay ID | Dye | Catalog # |
|---|---|---|---|
| ***ACTB*** | Hs01060665_g1 | FAM-MGB | 4331182 |
| ***ENTPD2*** | Hs00993192_g1 | | |
| ***TAS1R1*** | Hs01547926_g1 | | |
| ***TAS1R2*** | Hs01027711_m1 | | |
| ***TAS1R3*** | Hs00877447_g1 | | |
| ***TAS2R38*** | Hs00604294_s1 | | |
| ***POU2F3*** | Hs00205009_m1 | | |
| ***TRPM5*** | Hs00175822_m1 | | |
| ***PLCB2*** | Hs01080541_m1 | | |
| ***SNAP25*** | Hs00938957_m1 | | |
| ***CA4*** | Hs00426343_m1 | | |

Table S3. SYBR Green primers used for qPCR, related to Figure S2.

| Gene name | Forward | Reverse |
|---|---|---|
| ***ACTB*** | GAGCACAGAGCCTCGCCTTT | TCATCATCCATGGTGAGCTGG |
| ***AXIN2*** | GAGTGGACTTGTGCCGACTTCA | GGTGGCTGGTGCAAAGACATAG |
| ***SP5*** | CAGGCCTTTCTCCAGGACC | CGATGCGGCTACAGGTGG |
| ***NKD1*** | GGCAGCGGAGATGAGAAGAA | CGCACTGGAGCTCTTCAAAC |
| ***NOTUM*** | AGCAGTATCGCCACACAGAC | ACCCCGTTCCAGTACCTGAT |
| ***RNF43*** | GGTGAGTGGCCAGACTCAG | AAATGACCCGTAGCTCCTGC |
| ***ZNRF3*** | GCGACGCAGTCAGAATTCCA | CCCCTTGCTCTTGGAGTTGA |
| ***EPHB3*** | CACAAGTGAGAGAGGCTCTGG | CGAAGACAAGCCCAGCTGTA |
